# Revealing Protein-Level Functional Redundancy in the Human Gut Microbiome using Ultra-deep Metaproteomics

**DOI:** 10.1101/2021.07.15.452564

**Authors:** Leyuan Li, Zhibin Ning, Xu Zhang, James Butcher, Caitlin Simopoulos, Janice Mayne, Alain Stintzi, David R. Mack, Yang-Yu Liu, Daniel Figeys

## Abstract

Functional redundancy is a key property of ecosystems and represents the fact that phylogenetically unrelated taxa can play similar functional roles within an ecosystem. The redundancy of potential functions of human microbiome has been recently quantified using metagenomics data. Yet, the redundancy of functions which are actually expressed within the human microbiome remains largely unexplored. Here, we quantify the protein-level functional redundancy in the human gut microbiome using metaproteomics and network approaches. In particular, our ultra-deep metaproteomics approach revealed high protein-level functional redundancy and high nestedness in proteomic content networks - bipartite graphs that connect taxa with their expressed functions. We further examined multiple metaproteomics datasets and showed that various environmental factors, including individuality, biogeography, xenobiotics, and disease, significantly altered the protein-level functional redundancy. Finally, by projecting the bipartite proteomic content networks into unipartite weighted genus networks, functional hub genera across individual microbiomes were discovered, suggesting that there may be a universal principle of functional organization in microbiome assembly.

**Highlights:**

- Ultra-deep metaproteomics reveals high protein-level functional redundancy in the human gut microbiome
- Within-sample proteomic content networks display universal topology
- Various environmental factors influence the redundancy of expressed functions
- Functional hub genera are present across different datasets

## Introduction

The human gut microbiome is a complex ecosystem harboring trillions of microorganisms. Its taxonomic composition, functional activity and ecosystem processes have important consequences on human health and disease. It is crucial to study the human gut microbiome in the context of ecological communities (Gilbert and Lynch, 2019). Currently, the organizational principles and ecosystem functioning of the human gut microbiome remain under-investigated.

Structure-function relationships are a determining factor of ecological properties in the human gut microbiome (Vieira-Silva et al., 2016). Functional redundancy (FR) is considered to be one of the key ecological properties in microbial communities (Loreau, 2004). As a classical notion in community ecology, FR is closely related to the concept of ecological guilds, stating that species are grouped together based on the similarities of how they function in the community (Root, 1967; Wu et al., 2021). A high level of FR implies that members in a community maybe substitutable with little impact on the overall ecosystem functionality (Lawton and Brown, 1993).

Recently, a quantitative measure of FR for microbiome samples based on metagenomics data and the notion of genomic content network (GCN) was proposed (Tian et al., 2020). It was reported that the high level of FR in the human gut microbiome is related to a few topological features (e.g., the highly nested structure) of the GCN. Importantly, by definition, GCN-based FR calculations are derived from measuring the gene composition of a microbiome, without any regard for whether these genes are actually expressed. In other words, the within-sample FR calculated from the GCN only represents the redundancy of *potential* functions of a microbiome sample (i.e., the DNA-level FR), rather than the redundancy of actually *expressed* functions (e.g., protein-level FR).

Here we seek to quantify the redundancy of expressed functions for microbiome samples. To achieve that, functional microbiome profiling that can universally capture the expressed functions from microbes is required. Metaproteomics is a powerful tool that can bring microbiome studies to a level permissible to measuring expressed functions (Kleiner, 2019; Li and Figeys, 2020; Salvato et al., 2021). It identifies proteins and quantifies their abundances from microbiome samples based on liquid chromatography-tandem mass spectrometry (LC-MS/MS) techniques. In the last few years, metaproteomics has experienced an exponential growth in its identification coverage (Zhang et al., 2017), providing invaluable deep insights into the expressed functional activities of microbiomes. In a typical metaproteomics analysis, quantified proteins are used to determine the functional composition, taxonomic origins of the expressed functions, and to identify functional pathway changes through multivariate statistical analysis (Salvato et al., 2021; Zhang and Figeys, 2019).

In this study, we quantified the redundancy of expressed functions in the human gut microbiome based on ultra-deep metaproteomics data. Following the GCN approach (Tian et al., 2020), we constructed a proteomic content networks (PCN) for each microbiome sample by linking the taxa (identified from metaproteomics data) to their expressed proteins. Using tools from network science, we investigated the topological properties of these PCNs, and compared their matched GCNs from shotgun metagenomics. We next examined whether the topological properties of the PCNs are similar in metaproteomics datasets obtained by different analytical workflows and instrument platforms. Finally, we computed the associations between the protein-level FR and various host factors such as disease status and xenobiotics stimulation.

## Results

### Proteomic content networks are highly nested

We first constructed sample-specific PCNs using a dataset generated by an in-depth metaproteomics approach. Briefly, aliquots from four individuals’ ascending colon microbiome samples were subjected to protein extraction and digestion, followed by a high-pH reversed-phase fractionation (Batth et al., 2014) and a LC-MS/MS analysis (**Figure 1**). Metaproteomics RAW files were analyzed by our MetaPro-IQ approach (Zhang et al., 2016) using the integrated gene catalog (IGC) database of the human gut microbiome (Li et al., 2014). On average, 69,280 unique peptides and 30,686 protein groups were quantified per sample. The depth of peptide and protein quantification increased 54% and 49%, respectively, compared to our previously reported deep metaproteomics approach (Zhang et al., 2017) (**Supplementary Figure S1**). Using a “protein-peptide bridge” method (**Figure 1** and **Supplementary Notes**), functions that were annotated by protein groups and taxonomy that were identified by unique peptides were linked to construct the sample-specific PCN. These four microbiome samples were previously analyzed by metagenomics in the MetaPro-IQ study (Zhang et al., 2016). The metagenomics data were searched against the IGC database to construct the GCNs (see **Methods**). The PCNs achieved a reasonable depth and correlationship with the corresponding GCNs (**Supplementary Figure S2**).

**Figure 1.**
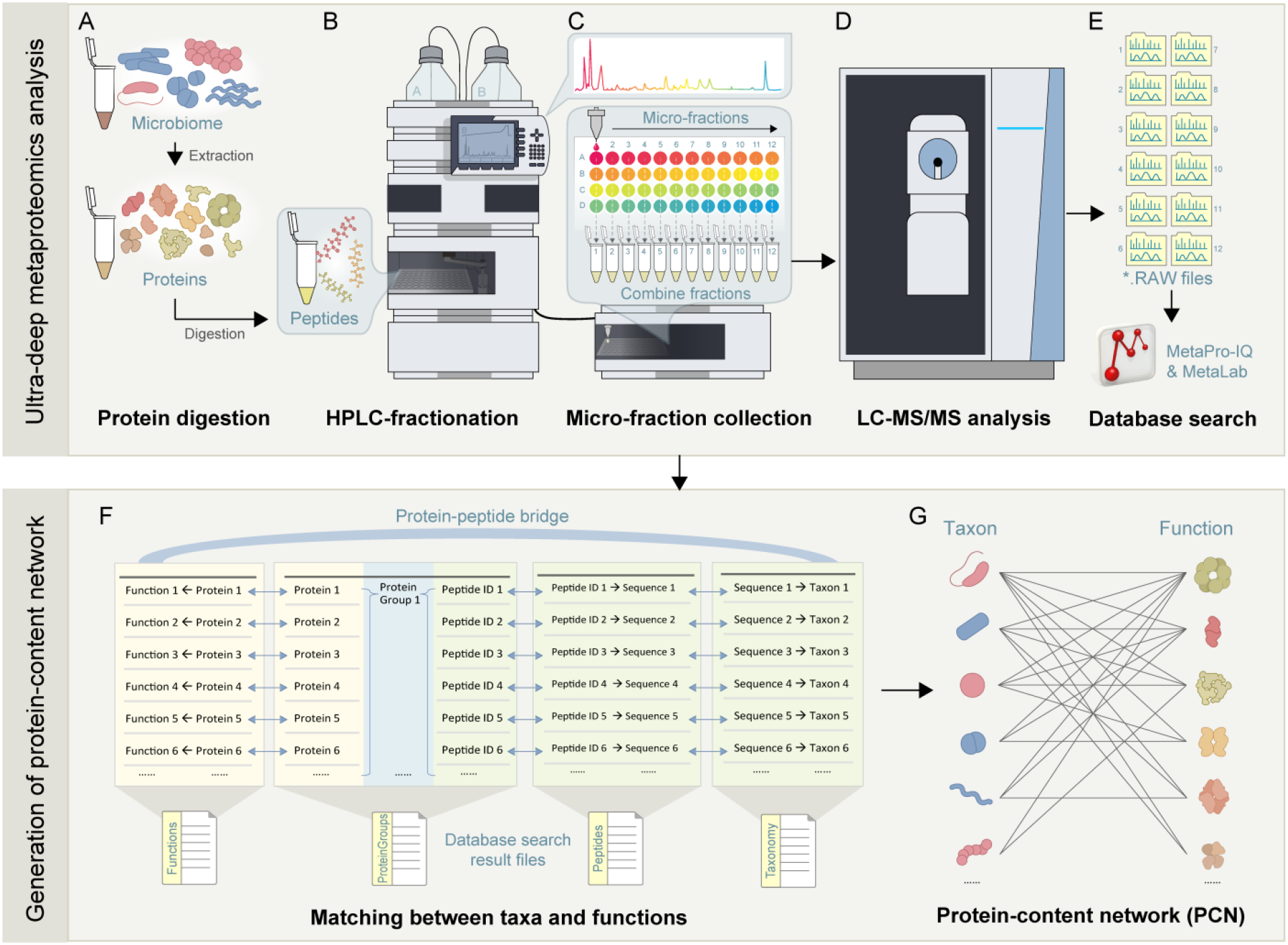
Generation of proteomic content network (PCN) using ultra-deep metaproteomics. A. Each individual’s gut microbiome sample was subjected to protein extraction. Then, purified proteins were digested by trypsin. B. The resulting peptides were fractionated using a high-pH reversed-phase approach. C. 48 micro-fractions were combined into 12 samples prior to LC-MS/MS analysis (D.). E. The LC-MS/MS *.RAW files were searched against the IGC database using MetaPro-IQ workflow and MetaLab. F. A protein-peptide bridge approach was used for generating the PCN (G.) from metaproteomics database search result files (see Methods and Supplementary Note).

We emphasize that those sample-specific GCNs can be combined to form a reference GCN for any given population, because the genomic content of a taxon should not be sample-dependent (Tian et al., 2020). By contrast, sample-specific PCNs cannot be combined to form a reference PCN, because the expressed protein content of any taxon is context-dependent. In that light, here we compare the GCN and PCN of each sample separately (**Figure 2**). **Figure 2B** shows a tripartite plot connecting microbial phyla and functional categories indicated from gene and proteins from one individual microbiome (HM454). This demonstrated that while some functional categories (e.g., energy production and conversion (C), carbohydrate metabolism and transport (G) etc.) showed expression from predicted functions in most phyla, there are functions (e.g., RNA processing and modification (A), mobilome (X) etc.) that were rarely expressed from the genes. Similar results were found for other samples (**Supplementary Figures S4-S6**). Due to the fact that some protein or peptide sequences are shared between two or more organisms in complex microbial communities, as a trade-off between depth and coverage, we analyzed the PCN at the genus level. The PCNs of individual microbiomes showed highly nested structures (**Figure 2A**). Nestedness metric based on Overlap and Decreasing Fill (NODF) showed that the PCNs in the four individuals’ microbiomes are highly nested networks (NODF = 0.42 ± 0.01, Mean ± SD, N = 4).

**Figure 2.**
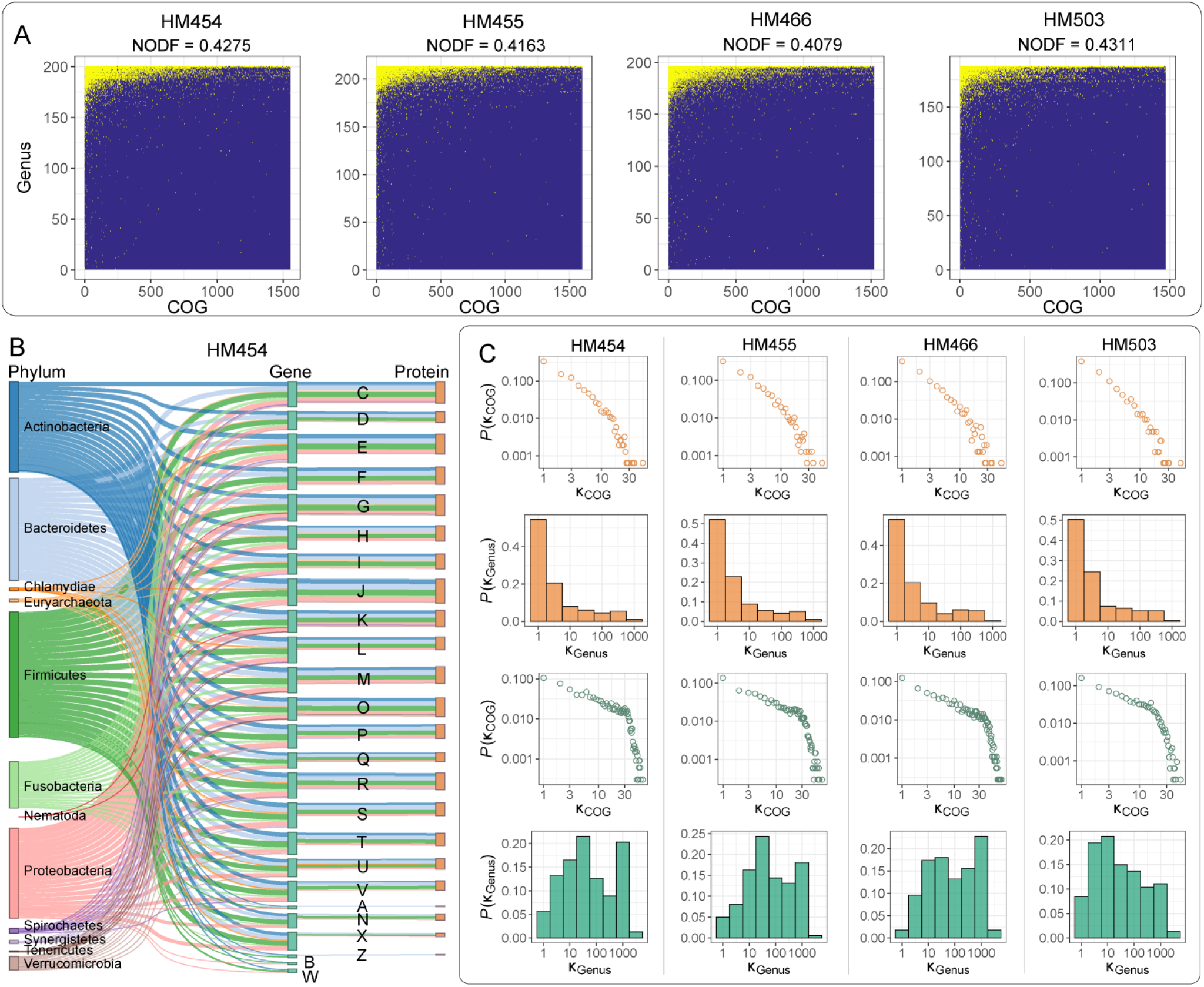
Proteomic content networks (PCNs) and genomic-content networks (GCNs) of individual microbiomes. A. Taxon-function incidence matrix of the PCN at the genus-COG level in the four individual microbiome samples. Here we used the classical Nestedness metric based on Overlap and Decreasing Fill (NODF) to characterize and visualize the nested structures of the bipartite taxon-function network, as described previously (Tian et al., 2020). The presences of genus-COG connections were shown in yellow points. B. A tripartite plot showing taxonomic and functional relationships between GCN and PCN in individual sample HM454. Letters represent different functional categories in the Clusters of Orthologous Groups (COGs) database. Similar results of the other three individual microbiomes are shown in **Supplementary Figure S4-S6.** C. The unweighted degree distribution of COGs in PCNs (first row), that of genera in PCNs (second row), that of COGs in GCNs (third row), and that of genera in GCNs (fourth row) in the four individual microbiomes.

We then calculated the degree distributions of genera and COGs in the PCNs and the GCNs, respectively. On the functional dimension, similar to previous observations in GCNs (Tian et al., 2020), the degree distributions of COGs in both the GCN and PCN have fat tails, representing COGs associated with a high number of taxa (**Figure 2C**). We performed enrichment analyses using the top-200 most linked COG nodes in the PCN incidence matrices, and discovered highly enriched functional categories of carbohydrate, amino acid and nucleotide metabolism, as well as energy production and conversion (**Supplementary Figure S3**). On the taxonomic dimension, in comparison to the PCN, the GCN had a relatively larger number of genera with high degrees (Figure 2C). This could be due to the fact that although many genera encoded genes for complex functional capacities, only a small subgroup were actually expressing these genes and thus actually playing complex functional roles.

### Redundancy differs between potential and expressed functions

We next compared the within-sample FR calculated from the metagenome (functional potentials) and the metaproteome (expressed functions), denoted as FRg and FRp, respectively. Following the definition of within-sample FR as previously described (Tian et al., 2020), we have

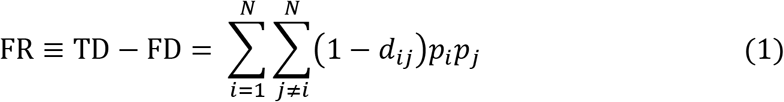

where 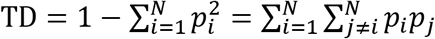 is the taxonomic diversity measured by the Gini-Simpson index, 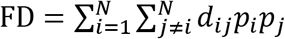 is the functional diversity measured by Rao’s quadratic entropy, *p_i_* is the relative abundance of taxon *i* in a community/sample of *N* taxa, *d_ij_* denotes the functional distance between taxa *i* and *j* measured by the weighted Jaccard distance between their genomic (or proteomic) contents (see **Methods**).

We emphasize that FR of a microbiome sample, as defined in Eq.(1), can be interpreted as the average functional similarity (or overlap) of two randomly chosen members in the sample. Since a potential function of any member in the microbiome sample may or may not be expressed under a certain environmental condition, we anticipate that the protein-level FR (i.e., FR_p_) of any microbiome sample should be no greater than its DNA-level FR (i.e., FR_g_). Indeed, as shown in **Figure 3A-B**, for each of the four individual microbiomes, we have FR_p_ < FR_g_ and nFR_p_ < nFR_g_, where nFR = FR/TD is the normalized FR. Interestingly, the FD (or TD) calculated from metagenomics and metaproteomics were comparable to each other (**Figure 3C-D**).

**Figure 3.**
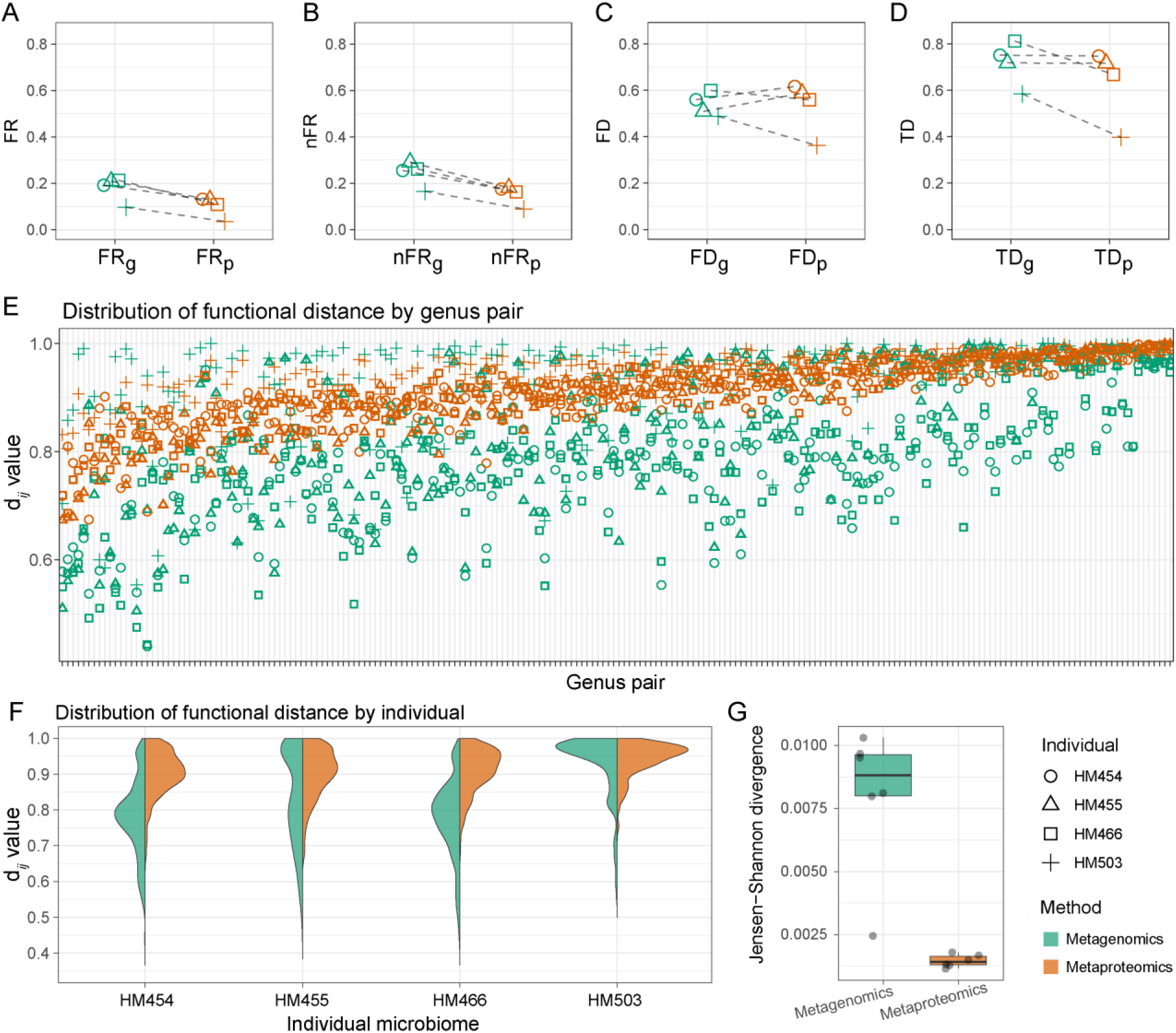
Redundancies of expressed functions and functional potentials. A. Within-sample functional redundancy (FR) in the metagenomes versus in the metaproteomes of the individual microbiomes. B. Within-sample FR normalized by taxonomic diversity (nFR) in the metagenomes versus in the metaproteomes of the individual microbiomes. C. Functional diversity (FD) in the metagenomes versus in the metaproteomes. D. Taxonomic diversity (TD) in the metagenomes versus in the metaproteomes. E. Functional distance (d*_ij_* value) between different pairs of genera in the metagenomes versus in the metaproteomes. F. Distribution of functional distance in the metagenomes versus in the metaproteomes in each individual microbiome. G. Pairwise comparisons of *d_ij_* distributions between individual microbiomes using Jensen-Shannon divergence. D-F were compared based on microbial genera of the top 95% overall protein biomass in the dataset.

Next, we investigated the functional distance *dij* between different taxa. For a metaproteome (or metagenome), *d_ij_* ∈ [0, 1] represents the dissimilarity of expressed functions (or functional capacities) between taxon-*i* and taxon-*j*, respectively. We calculated the *d_ij_* values between those genera that contributed to 95% of the genus-level protein biomass in the dataset. Interestingly, *d_ij_* values in the metagenomes were highly variable among individual microbiomes. By contrast, in the metaproteomes, *d_ij_* values between genera were more consistent (**Figure 3E**). We then compared the *d_ij_* values by individual microbiomes. *d_ij_* in metagenomics and metaproteomics were not linearly related (*R_adj_*^2^ = 0.47 ± 0.10, Mean ± SD, N = 4; **Supplementary Figure S7**). It was evident that the *d_ij_* distributions in the metagenomes varied dramatically across samples, whereas in the metaproteomes the *d_ij_* distributions were similar (**Figure 3E**). To quantify the variations, we performed pairwise comparisons between *d_ij_* distributions in the samples using Jensen-Shannon divergence. Result shows that the *d_ij_* distributions in metaproteomes were much more similar across individuals than in the metagenomes (**Figure 3F**). The similarity of *d_ij_* distributions in individual metaproteomes suggested that gut environments play an important role in shaping the microbial functional activities in an organized manner.

### Conserved PCN topology across metaproteomics platforms

We wondered whether different metaproteomic approaches could recapitulate the network properties of gut microbiomes’ PCNs. Routine metaproteomic analysis are often performed without fractionation. In addition, samples are analyzed with different analytical protocols, different parameters and using different models of LC-MS/MS platforms, etc. Here, we compared the topological properties of PCNs in four of our previously published datasets, briefly referred to as SISPROT (Zhang et al., 2017), RapidAIM (Li et al., 2020b), Berberine (Li et al., 2020a) and IBD (Zhang et al., 2018a) datasets, respectively. These four datasets vary considerably in the metaproteomic approaches used and in the types of environmental factors (xenobiotics, biogeography, diseases status etc.) being interrogated (see details in **Supplementary Table S1**).

It was notable that identification depths of these four datasets vary markedly, from 5,612 protein groups and 4,345 peptides per sample (Berberine) to 44,955 unique peptides and 20,558 protein groups per sample (SISPRORT)(**Supplementary Table S1**). We found that PCNs in all the four datasets displayed very similar topological structures with our new deep metaproteomics dataset (see Figure 1), i.e., highly nested structure, and heterogeneous degree distributions of both taxa and functions (**Figure 4**).

**Figure 4.**
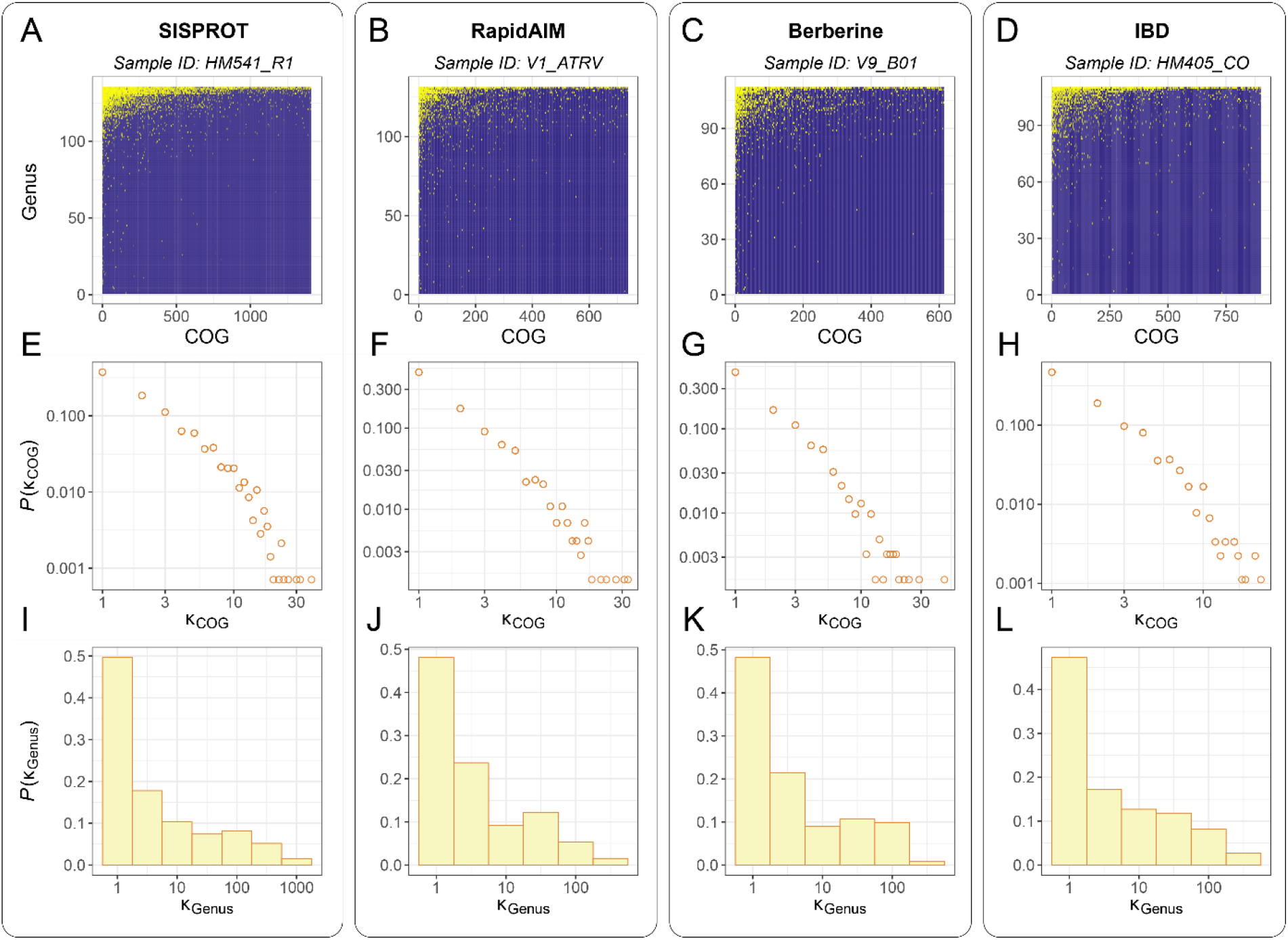
PCNs and corresponding degree distributions in different metaproteomics datasets. A-D. Taxon-function incidence matrix of the PCN corresponding to each metaproteomics dataset. The presences of genus-COG connections are shown as yellow dots. E-H. Unweighted degree distribution of COGs corresponding to each metaproteomics dataset. I-L. Unweighted degree distribution of genera corresponding to each metaproteomics dataset. Each vertical panel (gray-line box) represents the PCN of the first sample (by alphabet order) in each dataset. We also visualized the incidence matrices and degree distributions of all samples here: https://leyuan.shinyapps.io/pcn_visualization3/

### Redundancy of expressed functions is altered by environmental factors

Given that the PCN topological structures appeared to be universal across the four metaproteomic datasets, we can calculate and compare their protein-level functional redundancy FR_p_. The results showed that within-sample nFR_p_ values in these datasets were comparable to the previous deep metaproteomics data (**Figure 5A-D** versus **Figure 3A**). We performed within-dataset comparisons of nFR_p_ in response to different environmental factors. Significant inter-individual differences in nFR_p_ levels were observed (Wilcoxon rank-sum test; **Figure 5A-C**). In the RapidAIM and Berberine datasets, several xenobiotic compounds reduced nFR_p_ levels (**Figure 5E and G**). Among which, the antibiotic rifaximin showed the most impact on the individual microbiomes with nFR_p_ values decreasing 22.5 ± 9.4% (Mean ± SD, N = 5). Two-way ANOVA suggested that both inter-individual variation and effect of compounds significantly contributed to nFR_p_ variance (**Supplementary Tables S2-S3**). In patients diagnosed with inflammatory bowel disease (IBD), nFR_p_ levels were significantly lower than that of the non-IBD control individuals. There was no significant difference between the two different IBD subtypes Crohn’s disease (CD) and ulcerative colitis (UC) (**Figure 5D**). A significant decrease in nFR_p_ was found in inflamed regions from the terminal ileum (**Figure 5F**). Two-way ANOVA suggested significant contributions to nFR_p_ values by the diagnosis factor, as well as the inflammation factor which was nested in the biogeography factor (**Supplementary Table S4**).

**Figure 5.**
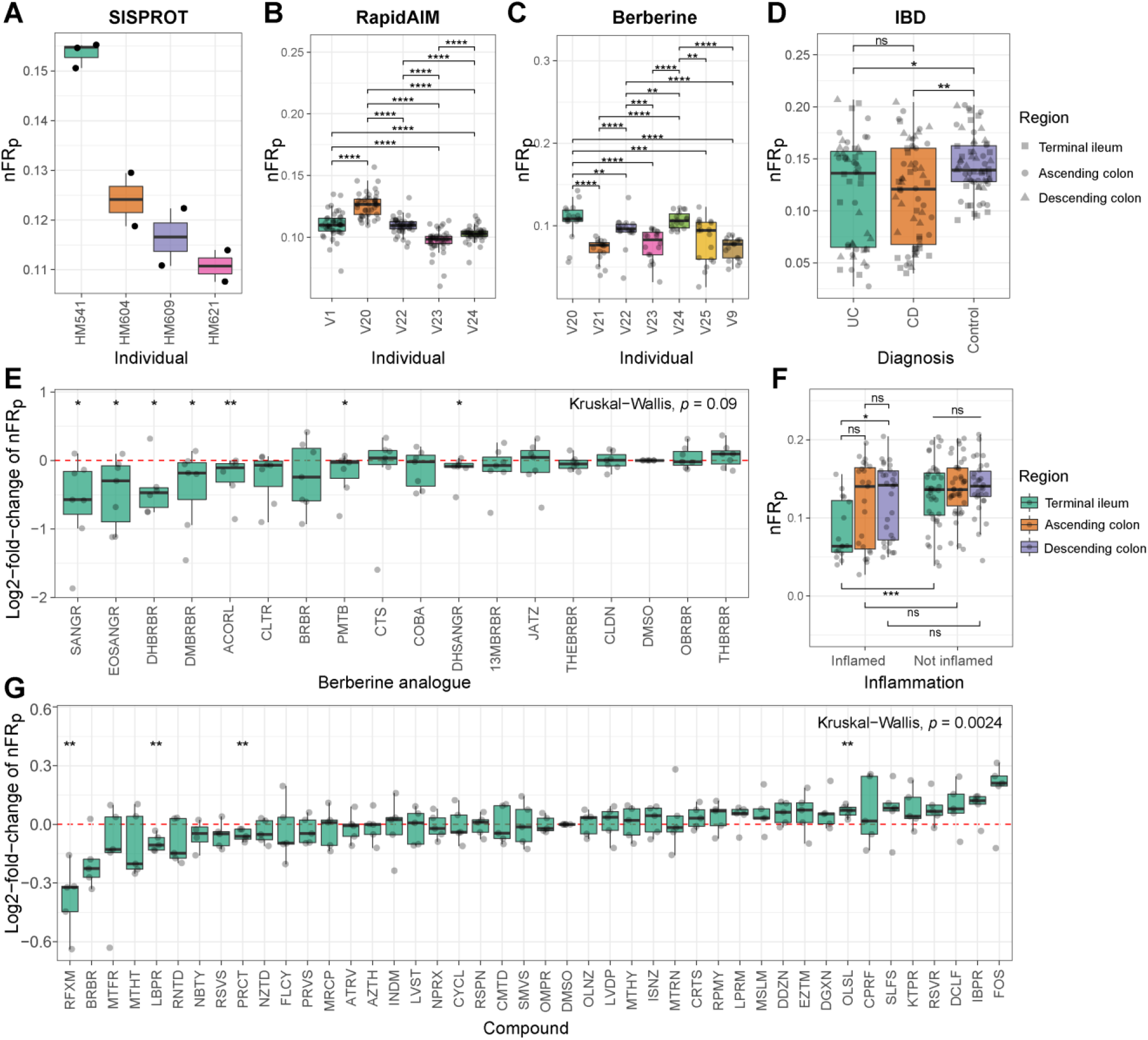
Functional redundancy comparisons in different metaproteomics datasets. A. nFR_p_ values by individual microbiomes in the SISPROT dataset. B. nFR_p_ values by individual microbiomes in the RapidAIM dataset. C. NODF values by individual microbiomes in the Berberine dataset. D. nFR_p_ values by diagnosis in the IBD dataset. E. nFR_p_ values by the presence of compounds in the Berberine dataset. F. nFR_p_ values by inflammation and gut region in the IBD dataset. G. nFR values by the presence of compounds in the RapidAIM dataset. Significance of differences between-groups were examined by Wilcoxon rank-sum test, *, **, *** and **** indicate statistical significance at the FDR-adjusted *p* < 0.05, 0.01, 0.001 and 0.0001 levels, respectively.

Despite global similarities of network properties across different individual samples and different metaproteomic approaches, we examined whether environmental factors had an impact on the nestedness of the PCNs. Similar to the nFR_p_ results, significant inter-individual differences in NODF values were observed (**Supplementary Figure S8 A-C**). Several compounds significantly decreased the NODF (**Supplementary Figure S8 E and G**). Patients diagnosed with IBD, as well as those with inflamed terminal ileums and/or ascending colons showed significantly decreased nestedness (**Supplementary Figure S8 D and F**). The agreement between within-sample nFR_p_ and NODF decrease in response to diseases and xenobiotic compounds further suggests that a nested topological structure is the key to high functional redundancy in a microbiome.

### Dissimilarity of functional expression between taxa is altered by xenobiotic compounds

To further elucidate the system-level functional mechanism behind the response of within-sample nFR_p_ to environmental alterations, we examined the metaproteomic functional distance d*_ij_* of different taxa. In the RapidAIM dataset, clear inter-individual differences could be found with principal component analysis (PCA) performed using d*_ij_* values (**Figure 6A**). Overall, under each drug treatment the d*_ij_* distributions appeared to be similar across the individual microbiomes (**Figure 6B**), and the d*_ij_* distribution (mean value across individual microbiomes, N=5) shifted upon treatment with several compounds as compared to the DMSO control (**Supplementary Figure S9**). Using a Permutation Multivariate Analysis of Variance (PERMANOVA) test, significant contributions from inter-individual difference, compound effects as well as individual-compound interactions were observed (*p* < 0.001; **Supplementary Table S5**). We quantified the dissimilarity of d*_ij_* distributions between drug treatments and the DMSO control using Kullback–Leibler (K-L) divergence. These results showed that ciprofloxacin, berberine, rifaximin, FOS, metronidazole, isoniazid, diclofenac and flucytosine significantly increased K-L divergence with the DMSO when compared to most other compounds (**Figure 6C**). This was in agreement with our previous findings that seven of these compounds (except flucytosine) resulted in global alterations in individual microbiome functionality (Li et al., 2020b). Similar findings were observed in the Berberine dataset, in which compounds that were previously found to alter microbiome functionalities (Li et al., 2020a) resulted in significant alterations in d*_ij_* distributions (**Supplementary Figure S10 and Supplementary Table S6**).

**Figure 6.**
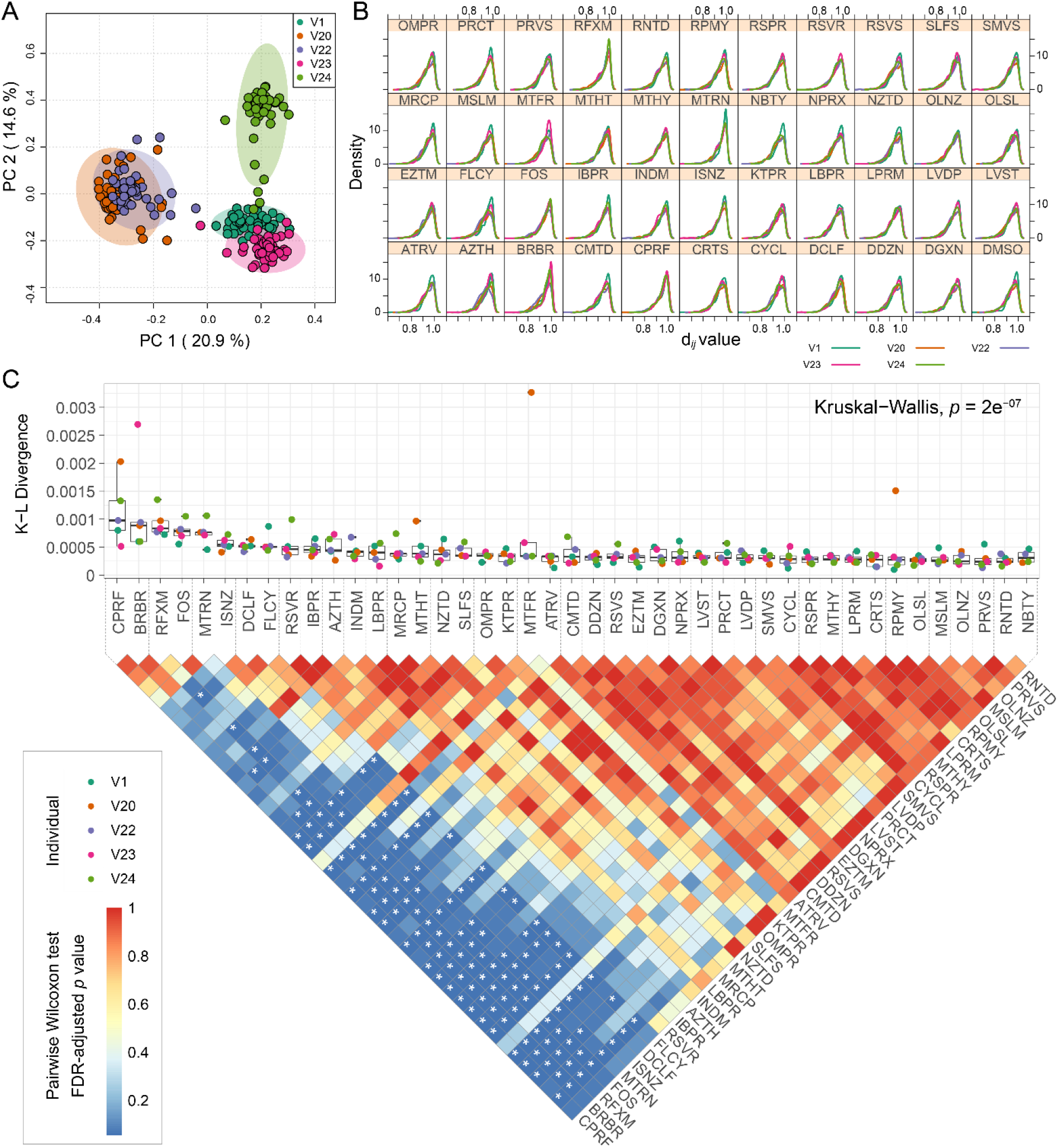
Between-genera functional distances in the RapidAIM dataset. A. Principal component analysis based on between-genera functional distances in individual metaproteomes. B. d*_ij_* distribution by the presence of different compounds and by different individual microbiomes. C. K-L divergence between the d*_ij_* distribution in the control (DMSO) and that of the other compounds. Kruskal-Wallis test result indicates that overall the compounds had heterogeneous levels of K-L divergence with the DMSO. Between-compound comparisons of the K-L divergence values were performed by a Pairwise Wilcoxon Rank Sum Tests, * indicates statistical significance at the FDR-adjusted *p* < 0.05 level. The results were based on microbial genera of the top 95% overall protein biomass in the dataset.

We further visualized the responses of d*_ij_* values using heatmaps (**Supplementary Figures S11** and **S12**). Certain genus pairs had similar d*_ij_* values under different drug treatments (in other words, consistently shown by a certain color range). By performing hierarchical clustering on the compound dimension, we observed different patterns of d*_ij_* responses. For example, in the RapidAIM dataset, antibiotics rifaximin, ciproflocaxin and metronidazole resulted in similar increases in functional distances between some pairs of genera. Several genera pairs e.g. *Prevotella* vs. *Subdoligranulum* and *Butyricicoccus* vs. *Clostridium* etc. showed larger functional distances compared to the cluster containing DMSO control; whereas genera pairs e.g. *Collinsella* vs. *Faecalibacterium* showed closer functional distance compared to other groups (**Supplementary Figure S11**). In the Berberine dataset, the d*_ij_* values between *Akkermansia* and a few other genera were increased by eight of the tested compounds (**Supplementary Figure S12**). We previously observed responses of *Akkermansia* to six of these compounds by differential protein analysis (Li et al., 2020a). However, response of *Akkermansia* in the presence of 6-ethoxysanguinarine (EOSANGR) and sanguinarine (SANGR) was not discovered before. This suggests that a system-level analysis of functional relationships between microbial taxa could sensitively provide a novel layer of information on microbial interrelations.

### Dissimilarity of functional expression between taxa is broadly increased by disease status

Similarly, we analyzed the metaproteomic functional distance between taxa using the IBD dataset. Clustering the data revealed that a subgroup of samples (the vertical cluster marked with red lines) showed an overall increase of d*_ij_* values between all visualized genera pairs (**Figure 7A**). These samples were mostly taken from the inflamed region of patients diagnosed with UC or CD. PERMANOVA test showed that d*_ij_* values differed significantly between diagnosed patients (especially inflamed regions) and the non-IBD controls (**Supplementary Table S7**). Overall, the d*_ij_* distributions in both UC and CD samples showed a rightward shift from the control samples (**Figure 7B**). Moreover, there was a rightward shift of the d*_ij_* distribution from healthy to inflamed gut regions (**Figure 7C**). These results explained why nFR_p_ was lower in diseased samples (**Figure 5D** and **F**). Interestingly, in a previous study based on function capacities inferred from 16S rRNA gene sequencing data, the d*_ij_* distribution (calculated at the OTU level) did not show a difference between IBD and control microbiomes (Tian et al., 2017). Volcano plot further showed that most of the d*_ij_* values between genera pairs were increased in the presence of inflammation (**Figure 7D**). This is different from compounds that affect specific pairs of genera in the microbiomes (described in the previous section) and suggests that inflammation disturbs the gut microbiomes’ functional organization by extensively weakening of the functional interrelations among microbes.

**Figure 7.**
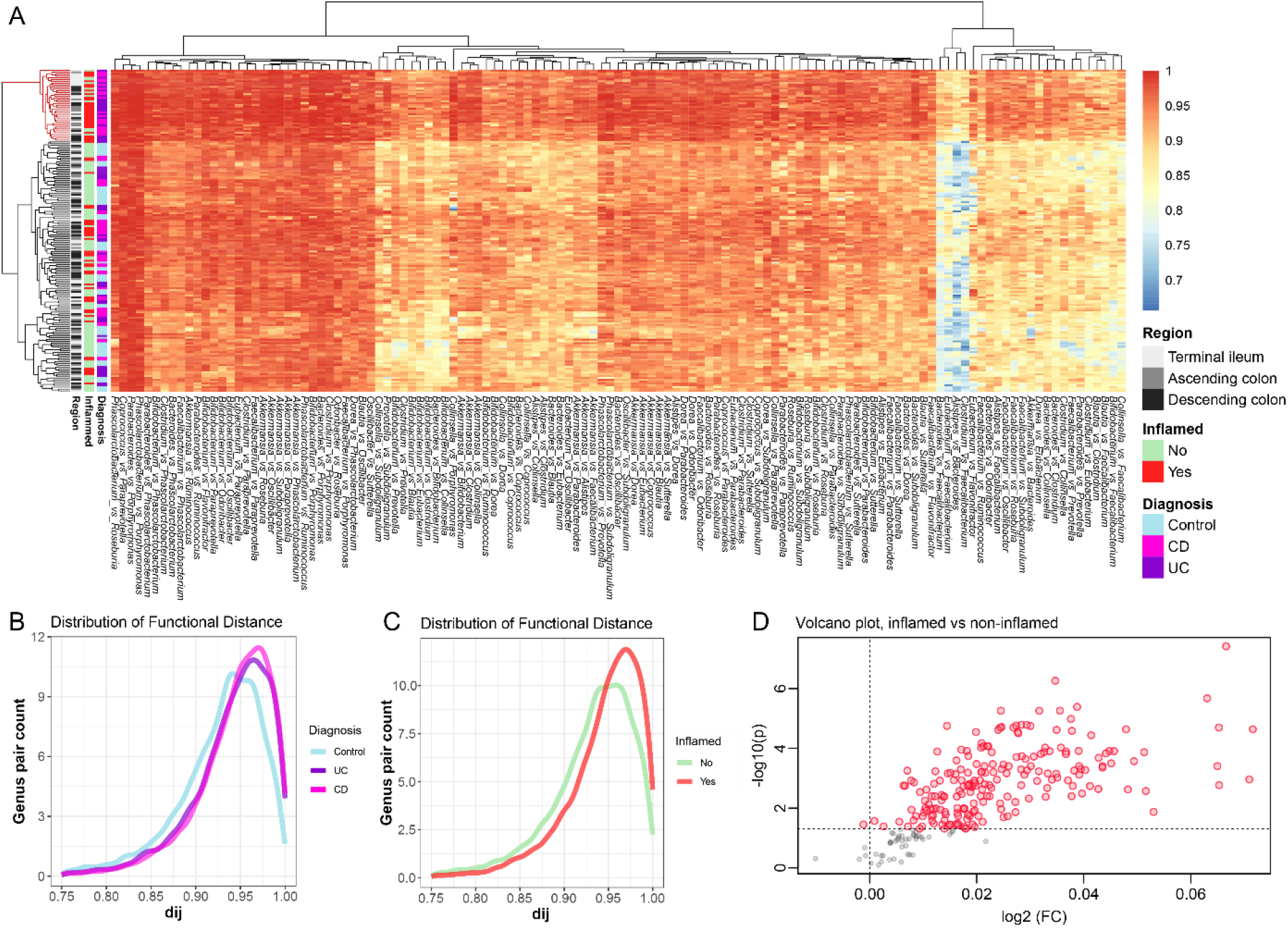
Between-genera functional distances in the IBD dataset. A. Heatmap showing d*_ij_* values between genera across samples in the IBD dataset. B. Distribution of d*_ij_* values by diagnosis. C. Distribution of d*_ij_* values by inflammation. D. Volcano plot showing altered d*_ij_* values between inflamed and non-inflamed sampling sites. The results were based on microbial genera of the top 95% overall protein biomass in the dataset.

### Global pattern of between-taxa functional association across datasets

Subsequently, we explored whether there was a universal pattern of functional interrelationships of protein expressions across individual gut microbiomes in our datasets. Thirteen abundant bacteria genera were consistently found in the five datasets (our in-depth dataset plus the four cross-platform datasets), i.e. *Bacteroides*, *Bifidobacterium*, *Blautia*, *Clostridium*, *Collinsella*, *Coprococcus*, *Dorea*, *Eubacterium*, *Faecalibacterium*, *Parabacteroides*, *Phascolarctobacterium*, *Roseburia* and *Ruminococcus*. We computed the functional distance (d*_ij_* values) between these genus pairs (**Figure 8A**) and used an Empirical Bayesian approach to correct for batch effects across platforms (**Supplementary Figure S13**). Box plots showed agreement of between-genera d*_ij_* values across all datasets (**Figure 8A**). Based on mean values of the functional distances (d*_ij_* cutoff >0.90), we constructed a global unipartite network of functional interrelations between microbial genera across the datasets (**Figure 8B**). *Eubacterium*, *Faecalibacterium*, *Ruminococcus*, *Bacteroides*, *Clostridium* and *Coprococcus* showed high number of linkages, suggesting that these “global hub genera” may play their roles as functional hubs in microbiomes. To validate this finding, we analyzed samples from another individual microbiome (not included in the above datasets) using our in-depth metaproteomics approach (**Supplementary Figure S1**). The functional interrelation network from this sample showed that most of our “global hub genera” had high degrees of connection in this new graph (**Figure 8C**).

**Figure 8.**
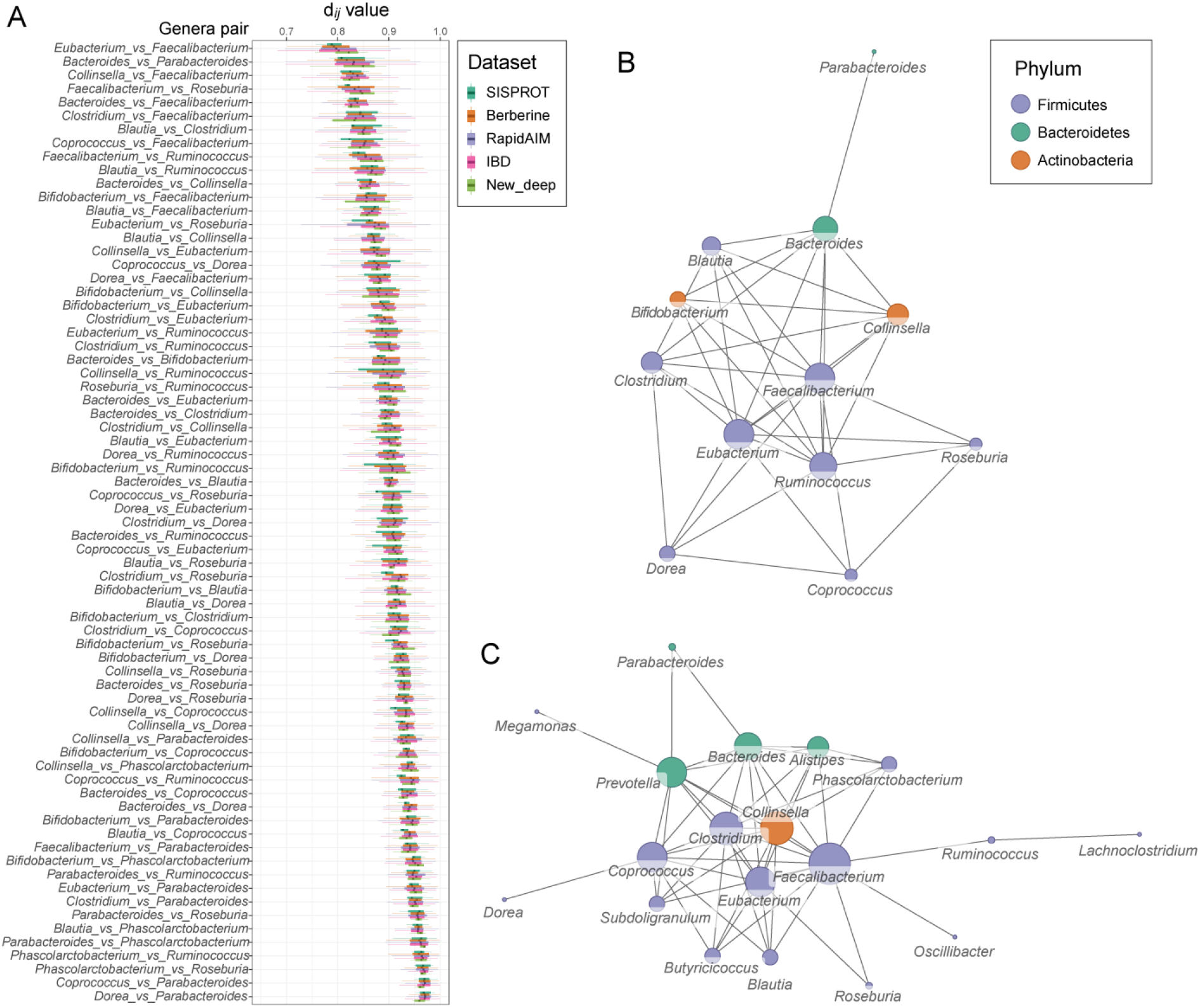
Comparison of between-genera *d_ij_* values across metaproteomics datasets. A. Comparison of d*_ij_* values across the five metaproteomics datasets in this study. Comparison was based on thirteen abundant microbial genera that were commonly found in these datasets. B. An average unipartite network projected from the taxon-function bipartite network based on the functional distances between microbial genera. A mean d*_ij_* values less than 0.9 across all data (N = 533) were shown as a linkage between two nodes. Size of a node corresponds to its degree. C. A unipartite network with an additional individual microbiome. The network was projected from the taxon-function bipartite network based on the functional distances between microbial genera. d*_ij_* value of 0.88 was used as the cut-off threshold on this graph. Size of a node corresponds to its degree.

## Discussion

A systems-oriented approach to understanding high-dimensional microbiome data can be employed by constructing of ecological networks (Angulo et al., 2019; Tian et al., 2020; Xiao et al., 2017). Network science (Barabási, 2013) provides a quantitative framework for representing and analyzing the principles underlying microbiome organization. Nevertheless, there has been a substantial gap between understanding microbiome assemblage and how its functionality is organized, which could not solely be examined by networks constructed from metagenomics. In this study, we demonstrated the usefulness of metaproteomics in gaining a system-level understanding of microbiome functionality by an in-depth investigation into the metaproteome network topology, functional redundancy and its contributing factors.

Using an in-depth metaproteomics approach, we showed that the human gut microbiome’s taxon-function networks at both the proteome and genome levels (i.e. PCN and GCN) are highly nested. In a microbiome PCN, the network being highly nested implies that specialist taxa tend to be playing functional roles that are a subset of active functions from generalist taxa (Bascompte and Jordano, 2007; Bascompte et al., 2003). Such functional network structures have been frequently found in macro-ecosystem networks of mutualistic interactions (food-webs) (Kondoh et al., 2010). Despite similarity in network topology between a microbiome’s metagenome and metaproteome, we found that the within-sample FR profiles differed markedly between expressed functions and functional capacities. The functional interrelationship of expressed proteomes between taxa appeared to be more robust across microbiomes, compared to that of the functional genomes. The COG degree distribution revealed that several functions were expressed in a high number of genera. These highly connected functions were enriched in metabolism of carbohydrates and amino acids, suggesting that microbial acquisition of nutrients from the environment and trophic interactions (Wang et al., 2019) between microbes could be major factors that shape their active functional organization. The nutrient-rich environment and mucosal immunity in the human gut provide a naturally selective growth condition for the microbes. Studies have shown that different host gut environments (human, mouse, rat and non-human primates) have distinct microbiome signatures (Nagpal et al., 2018; Nguyen et al., 2015). In human subjects, environmental factors such as diet and medication also significantly shape microbial community composition in the gut (Rothschild et al., 2018). Our result showing the robustness of between-taxa functional distances across individual microbiomes implies a more fundamental mechanism that underlies in the selective organization of microbiome functionalities by the environment.

Further, we found that taxon-function networks in metaproteomes showed universal properties: networks built with datasets generated by shallower metaproteomics approaches still capture the highly nested topology. This allowed us to make use of routinely generated metaproteomics datasets to observe the effects of multiple environmental factors, such as inter-individual variation, xenobiotics, disease and biogeography on the functional redundancy of the gut microbiome. We first showed that compounds with pharmacological activity can affect the redundancy of expressed functions in individual microbiomes. Overall distributions of functional distance between genera pairs were changed in response to some compounds, which was related to changes in a subset of between-genera functional distances. This suggests that xenobiotic compounds may affect functional redundancy by partially modifying the functional interrelationship between taxa.

Despite strong inter-individual signatures, we observed a universal pattern of between-taxa functional distances (d*_ij_*) across all analyzed datasets. Notably, this pattern was fully shifted by a global increase in d*_ij_* values and subsequently a significant decrease of the nFR in a subset of IBD samples mostly obtained from inflamed areas. Interestingly, this subset of samples still showed their own within-subset consistency in the distribution d*_ij_* values. This finding may support, from a functional angle, the hypothesis that there are alternative stable states (bi-stability or multi-stability) in the gut ecosystem(Gonze et al., 2017; Van de Guchte et al., 2020). One frequently discussed mechanism behind these alternative states has been the continuous exposure of the microbiome to a altered environmental parameter (Stein et al., 2013). An inflamed area in the gut will have a reduced mucus layer (van der Post et al., 2019) and elevated host defense responses(Zhang et al., 2018a). The host mucus layer is a nutritional source of cross-feeding in the gut microbiome(Bunesova et al., 2018; Kosciow and Deppenmeier, 2020; Schroeder, 2019). Loss of this layer may firstly affect the network hub functions of carbohydrate and amino acid metabolism, and subsequently affect the functional interactions in the whole community. In addition, host defense responses attenuate microbial oxidative stress responses(Zhang et al., 2018a), which has been associated to microbiome dysfunction (Luca et al., 2019). Decrease of within-sample FR has been associated with impaired microbiome stability and resilience(Moya and Ferrer, 2016). Resilient microbiota resist external pressures (e.g. antibiotics/dietary shifts) and return to their original state. Being non-resilient, a microbiome is likely to shift its composition permanently and stay at an altered new state instead of restoring to its original state of equilibrium(Dogra et al., 2020; Folke et al., 2004). Collectively, we disassembled the FR into one-to-one comparisons of between-taxa functional activities, and found that a global shift in functional roles of microbes towards a more heterogeneous pattern was a factor driving the decrease of FR and alteration of states in inflamed area in IBD patients.

Finally, the global pattern of between-genera functional distance across different metaproteomics datasets suggest that there may be universal community assemblage rules driven by the functional organization. In microbial community networks, highly linked nodes identified by a degree-based inference are often referred to as keystone taxa or hubs (Banerjee et al., 2018; Wang et al., 2017). Here, we refer such nodes observed in our PCNs as functional hubs. Across all metaproteomics datasets, *Eubacterium*, *Faecalibacterium*, *Ruminococcus*, *Bacteroides*, *Clostridium* and *Coprococcus* were found to be the most frequent functional hubs. Different approaches have been applied to identify keystone taxa in microbiomes with several agreements with our functional hubs(Fisher and Mehta, 2014; Ze et al., 2012). Such keystone taxa have been discussed as putative drivers of microbiome structure and function (Banerjee et al., 2018). Our current finding highlights the value of further investigation into functional hubs and hub functions in microbiome PCNs. This will provide a unique and systematic insight for the prediction of community functional responses, or for the manipulation of microbiome functioning.

## Acknowledgement

This work was partially funded by the Government of Canada through Genome Canada and the Ontario Genomics Institute (OGI-149) and the Ontario Ministry of Economic Development and Innovation (Project 13440). Y.-Y.L. acknowledges grants from National Institutes of Health (R01AI141529, R01HD093761, RF1AG067744, UH3OD023268, U19AI095219 and U01HL089856). The shotgun metagenomic analysis presented here was enabled in part by WestGrid (www.westgrid.ca) and Compute Canada (www.computecanada.ca). D.R.M. is partially supported through University of Ottawa Faculty of Medicine Distinguished Clinical Research Chair in Pediatric Inflammatory Bowel Disease. The authors acknowledge Ruth Singleton (Clinical Research Coordinator) for participant recruitment and data collection.

## Author contributions

Conceptualization, Y.-Y.L, D.F. and L.L.; Methodology, D.F., Y.-Y.L, L.L. and Z.N.; Formal Analysis: L.L.; Investigation L.L. and Z.N.; Resources, D.R.M., A.S., J.B., and J.M.; Data Curation: L.L., J.B., X.Z., Z.N. and C.S.; Writing –Original Draft, L.L., Y.-Y.L, and D.F.; Writing –Review & Editing, J.B., J.M. A.S., C.S., D.R.M., X.Z., and Z.N.; Visualization: L.L. and Z.N.; Supervision: D.F. and Y.-Y.L.

## Methods

**KEY RESOURCES TABLE.**

2 KEY RESOURCES TABLE
| Reagent or Resource | Source | Identifier |
| --- | --- | --- |
| **Biological samples** |  |  |
| ‘MetaPro-IQ’ samples | Lavage sample aliquots preserved at -80 °C | (Zhang et al., 2016) |
| ‘pepFunk’ samples | Cultured microbiome aliquots preserved at -80 °C | (Simopoulos et al., 2020) |
| **Metaproteomics reagents** |  |  |
| Ammonium formate | Sigma-Aldrich | 70221 |
| cOmplete™ mini tablet | Roche | C764L27 |
| Dithiothreitol | Sigma-Aldrich | 43815 |
| Formic acid | Sigma-Aldrich | F0507 |
| Hydrochloric acid | Fisher Chemical | A114S-500 |
| Iodoacetamide | Sigma-Aldrich | I1149 |
| Sep-Pak C18 1 cc Vac Cartridge | Waters | WAT054955 |
| TRIS (hydroxymethyl aminomethane) | OmniPur® | 9230 |
| Trypsin | Worthington Biochemical Corp., Lakewood, NJ | L5003740 |
| Urea | Sigma-Aldrich | U5378 |
| **Critical commercial assays** |  |  |
| DC™ protein assay reagents | Bio-Rad | 500-0113, 500-0114 and 500-0115 |
| **Software and algorithms** |  |  |
| R (versions 3.6.1 and 4.0.4) | R Foundation | http://www.r-project.org |
| Python (version 3.6) | Python | https://www.python.org |
| X!Tandem (version 2015.12.15.2) | X! Search Engine Development | https://www.thegpm.org/tandem/ |
| MaxQuant (version 1.5.2.8) | MaxQuant | https://www.maxquant.org |
| MetaLab (version 1.2.0) | iMetaLab Suite | https://imetalab.ca/ |
| DIAMOND (version 0.8.35) | DIAMOND | https://github.com/bbuchfink/diamond/releases/tag/v0.8.35 |
| MOCAT (version 1.3) | MOCAT | http://www.bork.embl.de/mocat/ |
| MEGAN (version 6.7.0) | MEGAN | https://www.wsi.uni-tuebingen.de/lehrstuehle/algorithms-in-bioinformatics/software/megan6/ |
| MetaPhlAn3 | MetaPhlAn | https://huttenhower.sph.harvard.edu/ |

### Resource availability

#### Lead Contact

Further information and requests for resources and reagents should be directed to the Lead Contact, Dr. Daniel Figeys.

### Materials availability

The study did not generate new unique reagents.

### Data and code availability

The ultra-deep metaproteomics datasets were deposited to the ProteomeXchange Consortium (http://www.proteomexchange.org) via the PRIDE partner repository. Database search outputs from the SISPROT (Zhang et al., 2017), RapidAIM(Li et al., 2020b), Berberine (Li et al., 2020a) and IBD (Zhang et al., 2018a) studies have been previously deposited to the ProteomeXchange Consortium with the dataset identifiers PXD005619, PXD012724, PXD015934 and PXD007819, respectively.

### Experimental subject details

Please refer to supplementary Table S1 in the MetaPro-IQ study (Zhang et al., 2016) for clinical details of the experimental subjects HM454, HM455, HM466 and HM503.

### Method details

#### Protein extraction and digestion

Protein extraction and digestion of the individual gut aspirate samples were performed as described previously(Zhang et al., 2018b), with minor modifications. Frozen aliquots of aspirate samples were thawed and subjected to differential centrifugation for microbial cell purification: the samples were first centrifuged at 300 g, 4 °C for 5 min to remove debris; the resulting supernatant was centrifuged at 14,000 g for 20 min to pellet microbial cells; the pellets were then washed three times by resuspending in cold phosphate-buffered saline (PBS) and centrifuging at 14,000 g, 4 °C for 20 min. Next, the washed microbial cell pellets were resuspended in a cell lysis buffer containing 4% sodium dodecyl sulfate (w/v), 8 M urea, 50 mM Tris-HCl (pH = 8.0), and one Roche cOmplete™ mini tablet per 10 mL buffer, followed by ultra-sonication (30 s on, 1 min off, amplitude of 25%, two rounds) using a Q125 Sonicator (Qsonica, LLC). Cell debris was then removed by a centrifugation at 16,000 g, 4 °C for 10 min.

Each of the protein extract was then precipitated in five times its volume of precipitation solution (acetone : ethanol : acetic acid = 49.5 : 49.5 : 1, v:v:v) at −20 °C overnight. The precipitated proteins were pelleted by centrifuging at 16,000 g, 4 °C for 20 min, followed by being washed with ice-cold acetone for three times to remove excess SDS that may affect trypsin activity. Next, the washed proteins were resuspended in a buffer containing 6M urea and 1M Tris-HCl (pH = 8.0). Protein concentration was determined by the DC™ assay (Bio-Rad Laboratories, Canada) following the manufacturer’s manual.

Finally, proteins were subjected to an in-solution tryptic digestion. The samples were reduced in 10 mM dithiothreitol (DTT) at 56 °C for 30 min, then were alkylated by 20 mM iodoacetamide (IAA) at room temperature in dark for 40 min. The samples were then diluted 10 times with 1 M Tris-HCl buffer (pH = 8.0), followed by trypsin digestion (at a concentration of 1 μg trypsin per 50 μg proteins) at 37 °C for 24 hours. The digests were then acidified to pH = 3 using 10% formic acid, followed by a desalting step using Sep-Pak C18 Cartridge (Waters, Milford, MA, USA). The cartridges were first activated using 100% acetonitrile, and then equilibrated using 0.1% formic acid (v/v) before passing samples through the columns for three times. Samples bonded to the cartridges were then washed using 0.1% formic acid (v/v), and finally the samples were eluted from the cartridges using the elution solution containing 80% acetonitrile and 0.1% formic acid (v/v).

### High-pH reversed phase fractionation

Eluted samples were evaporated in a SAVANT SPD1010 SpeedVac Concentrator (Thermo Fisher Scientific, USA), and resuspended in 0.1% formic acid (v/v) to a concentration of 1 μg/μL for high-pH reversed phase fractionation following a previous workflow(Batth et al., 2014), with minor adaptations: 30 μL sample were loaded to a ZORBAX Bonus-RP column (with 3.5 μm C18 resins, ID 2.1 mm, length 50 mm; Agilent Technologies, USA), and fractionated with a Agilent 1200 series HPLC System (Agilent Technologies, Germany). A 60-min gradient consisting of 5 - 35% buffer B (v/v) in 1 - 42 min, and 35 - 50% buffer B in 42 - 45 min at a flow rate of 100 μL/min was used for the fractionation. Here, 10 mM ammonium formate was used as buffer A, and 10 mM ammonium formate with 90% acetonitrile (v/v) was used as buffer B. Ammonium hydroxide was used to adjust the pH of both buffers A and B to 10. Sample fractions were continuously collected into 96 well plates by an Agilent 1100 Series Micro-FC G1364D micro fraction collector (Agilent Technologies, Germany). For each sample, 48 fractions were collected into different wells at 1 min intervals over the first 48 min. The samples were then pooled by combining four fractions at an interval of 12 wells, resulting in 12 fractionated samples per individual microbiome (**Figure 1A**).

### HPLC-ESI-MS/MS analysis

After evaporation and resuspension in 0.1% formic acid, each fraction was analysed by HPLC-ESI-MS/MS consisting of an UltiMate 3000 RSLCnano system (Thermo Fisher Scientific, USA) and an Orbitrap Exploris 480 mass spectrometer (Thermo Fisher Scientific, USA). A 60-min gradient of 5 to 35% (v/v) buffer B at a 300 μL/min flow rate was used to separate the peptides on a tip column (75 μm inner diameter × 10 cm) packed with reverse phase beads (3 μm/120 Å ReproSil-Pur C18 resin, Dr. Maisch GmbH, Ammerbuch, Germany). Here, 0.1% formic acid (v/v) was used as buffer A, and 0.1% formic acid with 80% acetonitrile (v/v) was used as buffer B. The MS full scan ranging from 350 – 1400 m/z was recorded in profile mode with the resolution of 60,000. Data-dependent MS/MS scan was performed with the 12 most intense ions with the resolution of 15,000. Dynamic exclusion was enabled for duration of 30 s with a repeat count of one.

### Database search

Database search for the fractionated metaproteomics samples was performed based on the MetaPro-IQ workflow(Zhang et al., 2016). Briefly, a two-step database search was first performed using X!Tandem (version 2015.12.15.2). All sample fractions were searched against the integrated gene catalog (IGC) of human gut microbiome (http://meta.genomics.cn/)(Li et al., 2014) to generate a reduced database, then a classical target-decoy database search was performed using the reduced database to generate confidently identified peptide and protein lists based on a strict filtering criteria of FDR = 0.01. The protein lists for all sample fractions were then combined, and duplicated proteins were removed to generate a combined non-redundant FASTA database using an in-house PERL script. Next, MaxQuant (version 1.5.2.8) was used to generate quantified protein groups and peptides in each sample using the combined non-redundant FASTA database. Carbamidomethylation of cysteine was set as a fixed modification, oxidation of methionine and N-terminal acetylation were set as potential modifications. The maximum missed cleavages of trypsin was set as two. The resulting peptide and protein group lists from MaxQuant were then inputted to MetaLab (version 1.2.0) for taxonomic analysis and functional annotation(Cheng et al., 2017). For the taxonomic analysis, identified peptides were mapped to taxonomic lineages based on a built-in pep2tax database in MetaLab. Functional annotation to COG was performed against a database generated by mapping proteins in the IGC database to clusters of orthologous groups (COGs) using Diamond (version0.8.35). The dataset was deposited to the ProteomeXchange Consortium (http://www.proteomexchange.org) via the PRIDE partner repository. We directly used MetaPro-IQ or MetaLab (which automates MetaPro-IQ) database search outputs from the SISPROT (Zhang et al., 2017), RapidAIM(Li et al., 2020b), Berberine (Li et al., 2020a) and IBD (Zhang et al., 2018a) studies. They have been previously deposited to the ProteomeXchange Consortium with the dataset identifiers PXD005619, PXD012724, PXD015934 and PXD007819, respectively.

For the metagenomics analysis, data were obtained from the previous MetaPro-IQ study (Zhang et al., 2016), accessible from the NCBI sequence read archive (SRA) under the accession of SRP068619. To enable the comparison between GCN and PCN, here we reanalysed the raw metagenomics reads by searching against the IGC database. First, the raw reads were processed using MOCAT for trimming and quality filtering, and for human reads removal as previously described (Zhang et al., 2016). Next, the cleaned/non-human paired end reads were used for DIAMOND against the IGC database. DIAMOND results of paired end reads were then merged, an annotation was confirmed only when both R1 and R2 were matched to the same protein or proteins. Finally, the result of each sample was summarized to generate a list of proteins and their corresponding read numbers.

### Metaproteomic and metagenomic content networks

For the generation of metaproteomic content networks (PCNs), a ‘peptide-protein bridge’ approach (see details in **Supplementary Note**) was used to match functions to taxa based on the database search output files, i.e. peptides, protein groups, taxonomy and function. The protein groups table (generated by MaxQuant) contains information on the identified proteins, and identifiers of peptide sequence associated to each protein group. The taxonomy table generated by MetaLab contains peptide sequences and their corresponding lowest common ancestor (LCA) taxa. The function table contains identified proteins and their corresponding functional annotations. Therefore, at we first matched the protein groups to taxa through the peptides. Next, functions of the proteins were combined to the list to generate a taxon-to-function table that was bridged by the peptide-protein identification relationship. Protein group intensity was used as the quantification information in PCNs. Then, a PCN of *N* taxa and *M* functions can then be represented by an *N* × *M* incidence matrix **P** = [P_ia_], where P_ia_ ≥ 0 is the total intensity of proteins of function-*a* in taxon-*i* normalized by the total intensities of functional proteins in taxon-*i*.

To generate GCNs from the IGC search result, the same functional annotation database as in MetaLab was used to annotate identified proteins to COGs. The taxonomic information of proteins was obtained by searching against an in-house database, which was generated by querying IGC proteins against the NCBI non-redundant (nr) database (downloaded 2/3/2016) using DIAMOND, and outputting the taxonomic lineages using MEGAN (version 6.7.0). The count of raw reads corresponding to each protein was used as the quantification information in GCNs. Similarly, the GCN can then be represented as **G** = [G_ia_], where G_ia_ ≥ 0 is the raw read counts of proteins of function-*a* in taxon-*i* normalized by the total counts of raw reads in taxon-*i*.

### Calculation of functional distance and functional redundancy

Weighted Jaccard distance *d_ij_* between metagenomic (or metaproteomic) contents of taxon-*i* and *j* can then be calculated with the GCN and PCN profiles **G** and **P**, respectively, as described previously (Tian et al., 2020). For GCN, we have

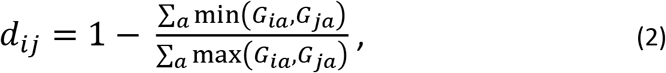

and for PCN, we have

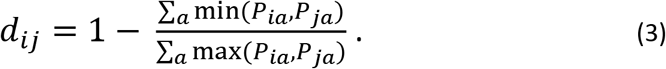

The relative abundance of taxon-*i* in each community was denoted as *p_i_*. In each metagenomics sample, *p_i_* was quantified using MetaPhlAn3 with default settings. In each metaproteomics sample, *p_i_* was quantified using the total abundances of unique peptides corresponding to taxon-*i*. With the *d_ij_* and *p_i_* values, within-sample FR of the metagenomic and the metaproteomic profiles, denoted as FR_g_ and FR_p_, respectively, were then calculated according to equation (1) given in the Results section.

### Statistical analysis and visualization

The statistical details of analysis can be found in the figure legends and in the main texts, including the statistical tests used and significance criteria. Computation of GCN, PCN and functional redundancy were performed using in-house Python codes. NODF values were computed using the R package RInSp. Jensen-Shannon divergence and Kullback–Leibler divergence were calculated using the R package LaplacesDemon. Two-way ANOVA was performed using R function aov(). PERMANOVA tests were performed using R packages “vegan” and “BiodiversityR”. Kruskal-Wallis and Wilcoxon rank sum tests were performed using R functions kruskal.test() and wilcox.test(), respectively. Network incidence matrices, degree distributions, bar plots, box plots, and violin plots were visualized using the R package ggplot2. Unipartite networks were visualized using the R package igraph. Tripartite networks were visualized using the R package networkD3. Heatmaps were visualized using the R package pheatmap. Volcano plot was analyzed by MetaboAnalyst (version 4.0) under non-parametric test setting. The interactive webpage (https://shiny2.imetalab.ca/shiny/rstudio/PCN_visualizer/) for visualization of all GCNs and PCNs analyzed in this paper was created using the R packages shiny and shinydashboard.

