## Supplementary figures for "Revealing Protein-Level Functional Redundancy in the Human Gut Microbiome using Ultra-deep Metaproteomics"

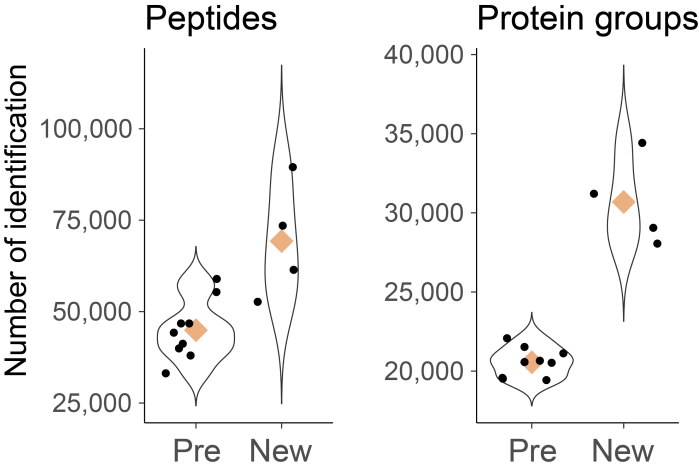

**Figure S1. Comparison of sequencing depth**

Comparison of sequencing depth between previously reported deep metaproteomics (Pre) dataset (Zhang *et al*., 2017) and the new deep metaproteomics approach (New).

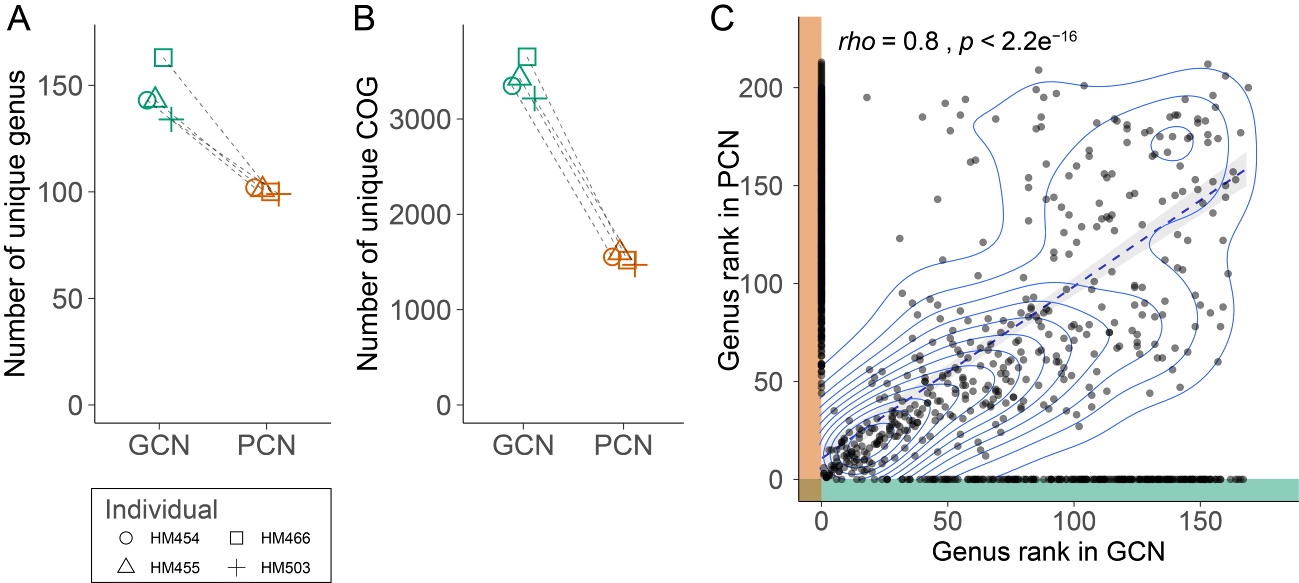

**Figure S2. Comparison of feature dimensions in GCN and PCN**

A-B. Comparison of genus- (A) and COG-level (B) identification between GCN and PCN. Number of genera that have at least 3 unique peptides was counted here for the PCN. C. Scatter plot comparing the rank of genus by number of identified COGs in GCN and PCN. *rho* and *p* values represent non-parametric Spearman correlation of the shared genera between GCN and PCN. Dashed line was the fitted linear model of the shared genera, and contour represents the density of the scatter points. The PCN in Panel C included the genera containing less than 3 unique peptides.

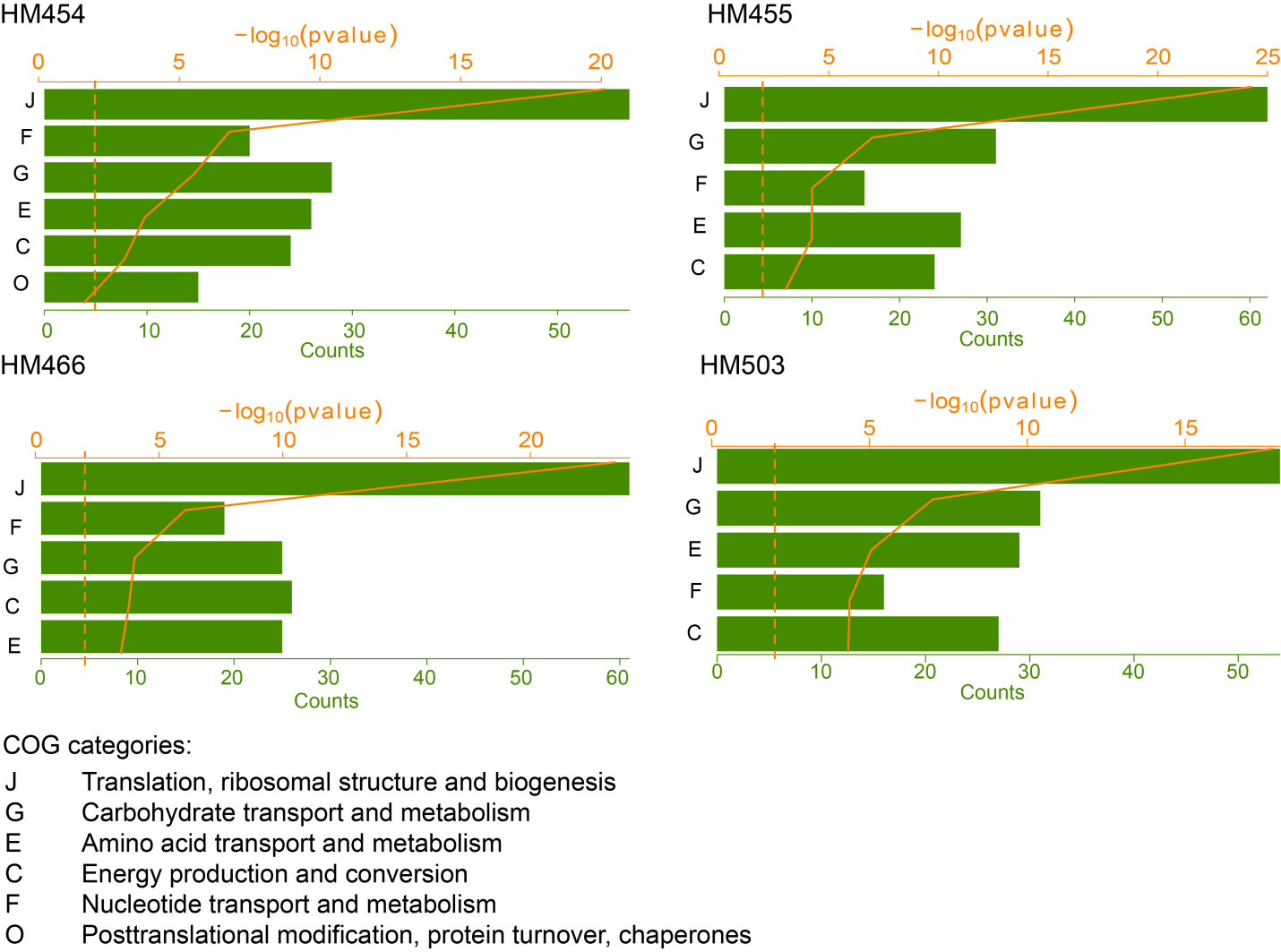

**Figure S3. Enrichment analysis of highly linked COG nodes.**

Top 200 COG accessions in each individual microbiome’s PCN incidence matrix were subjected to enrichment analysis using our online Shiny App “Enrichment Analysis” (<https://shiny.imetalab.ca/metaproteomics_enrichment/>). COG category was used as the enrichment analysis type, and *P*-adjusted to filter the enrichment was set at 0.05.

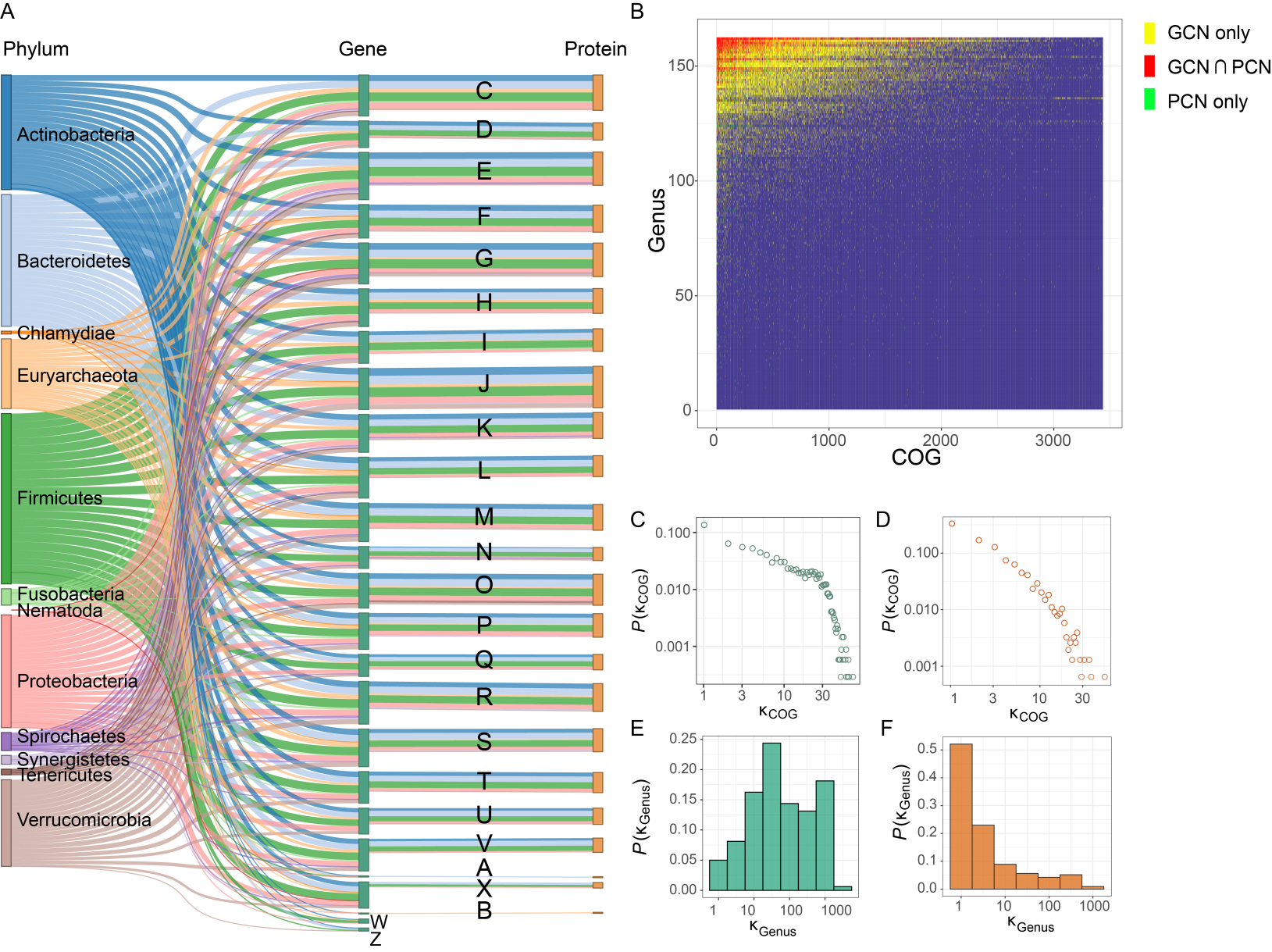

**Figure S4. A tripartite plot showing taxonomic and functional relationships between GCN and PCN for individual microbiome sample HM455.**

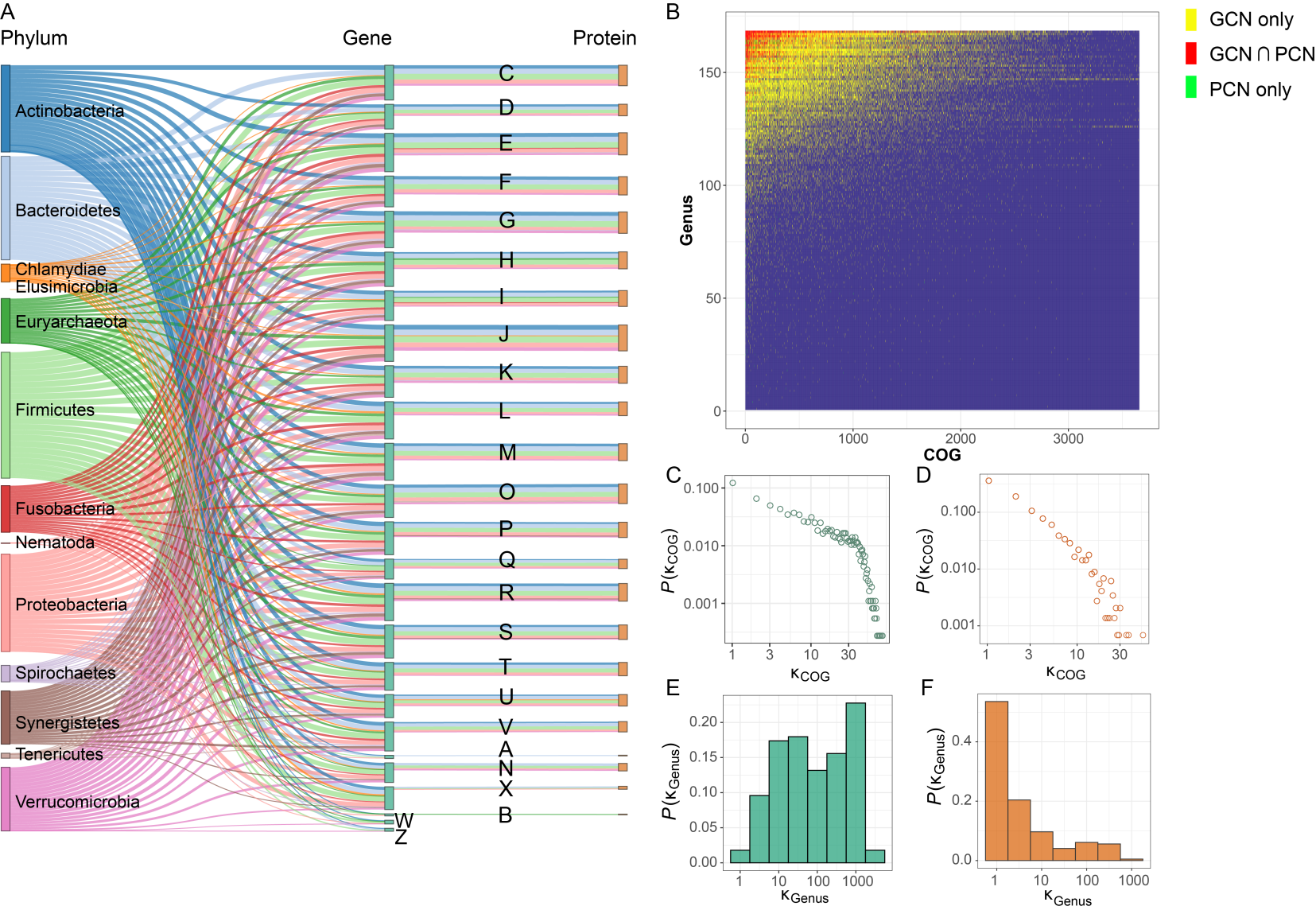

**Figure S5. A tripartite plot showing taxonomic and functional relationships between GCN and PCN for individual microbiome sample HM466.**

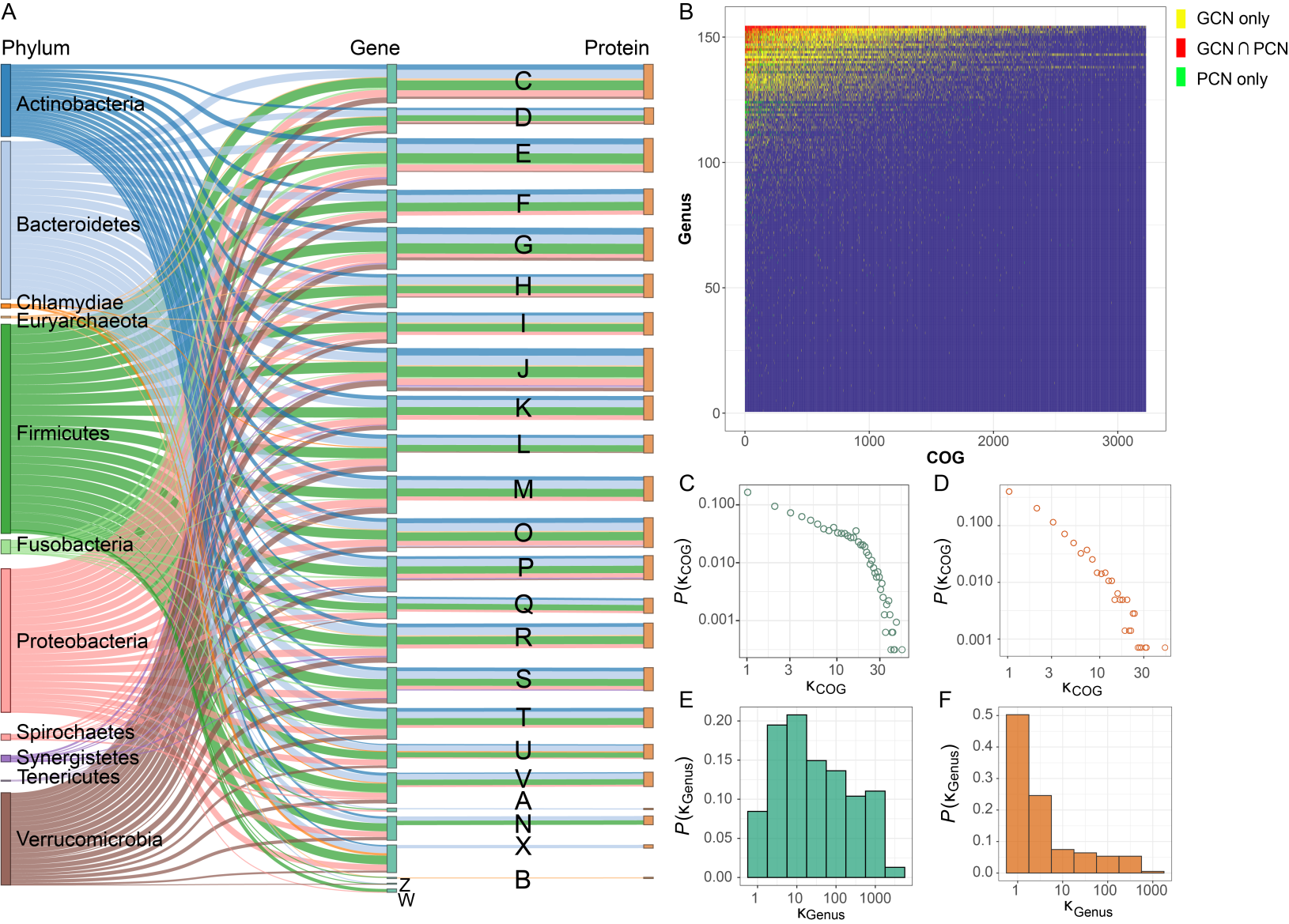

**Figure S6. A tripartite plot showing taxonomic and functional relationships between GCN and PCN for individual microbiome sample HM503.**

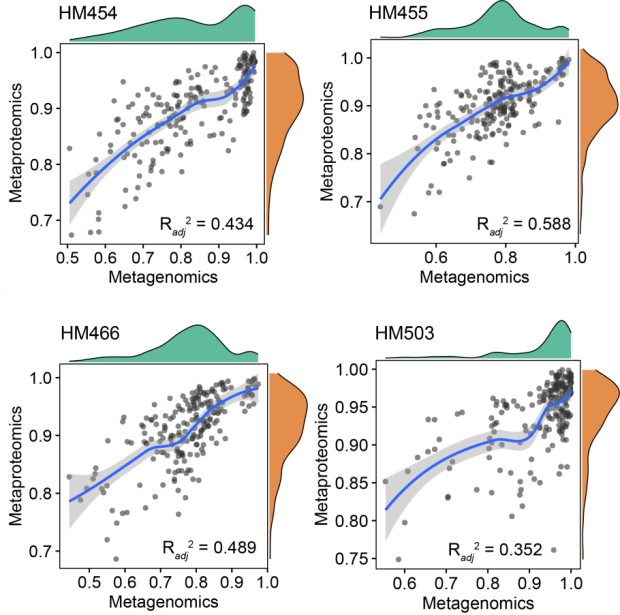

**Figure S7. Comparison of functional distance between metaproteomics and metagenomics**

Scatter plots compared the values between functional distances of genera pairs in the metagenomics and the metaproteomics data of individual microbiomes. Density plots represent the distribution of d*_ij_* values on the corresponding axis. Adjusted R^2^ value indicated the goodness of linear regression between metagenomics and metaproteomics d*_ij_* values in each individual microbiome. Microbial genera comprising the top 95% protein biomass were used in this figure.

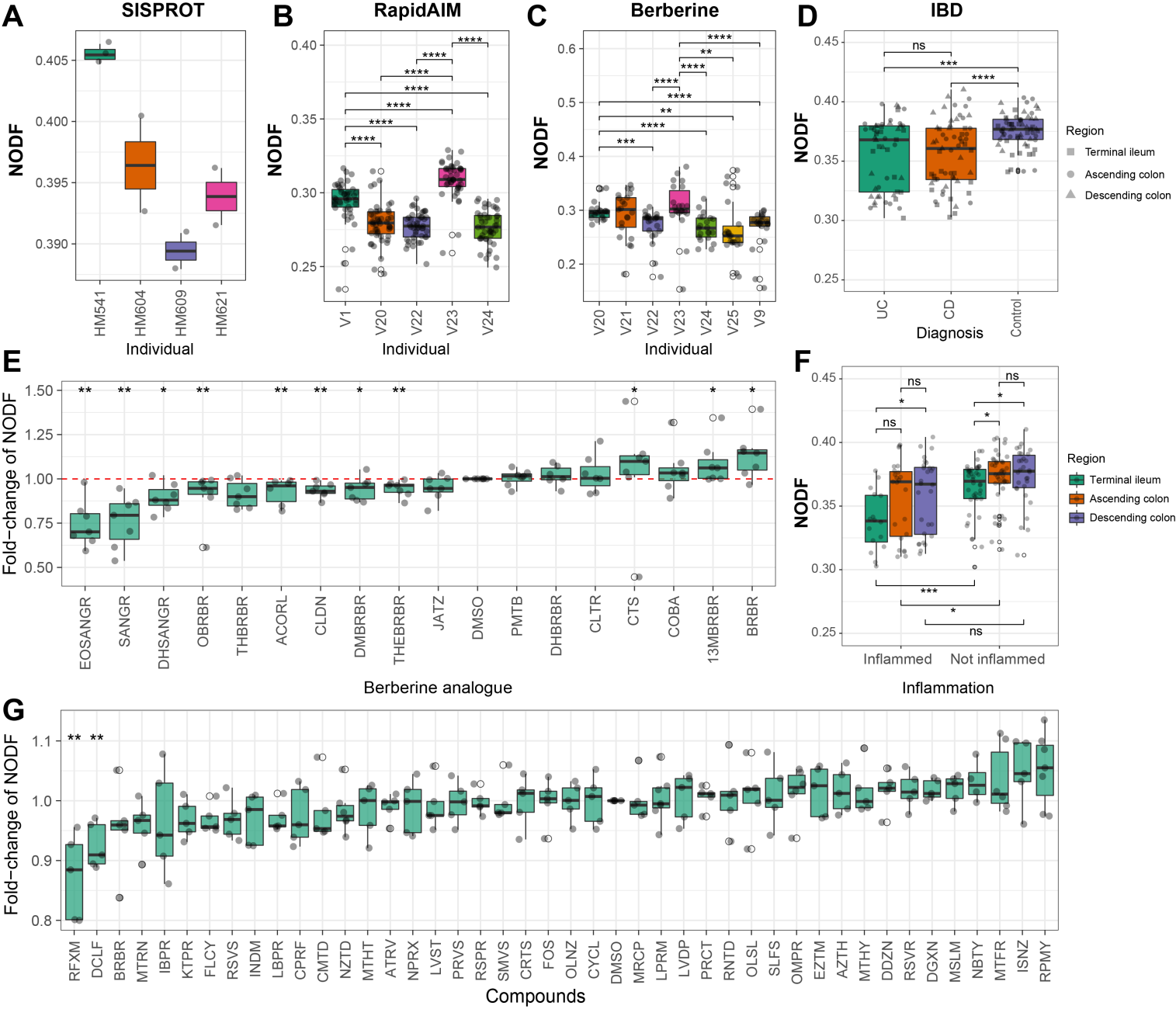

**Figure S8. Nestedness metric based on Overlap and Decreasing Fill (NODF) of metaproteomics datasets.** A. NODF values by individual microbiomes in the SISPROT dataset. B. NODF values by individual microbiomes in the RapidAIM dataset. C. NODF values by individual microbiomes in the Berberine dataset. D. NODF values by diagnosis in the IBD dataset. E. NODF values by the presence of compounds in the Berberine dataset. F. NODF values by inflammation and gut region in the IBD dataset. G. NODF values by the presence of compounds in the RapidAIM dataset. Significance of differences between-groups were examined by Wilcoxon rank-sum test, *, **, *** and **** indicate statistical significance at the FDR-adjusted *p* < 0.05, 0.01, 0.001 and 0.0001 levels, respectively.

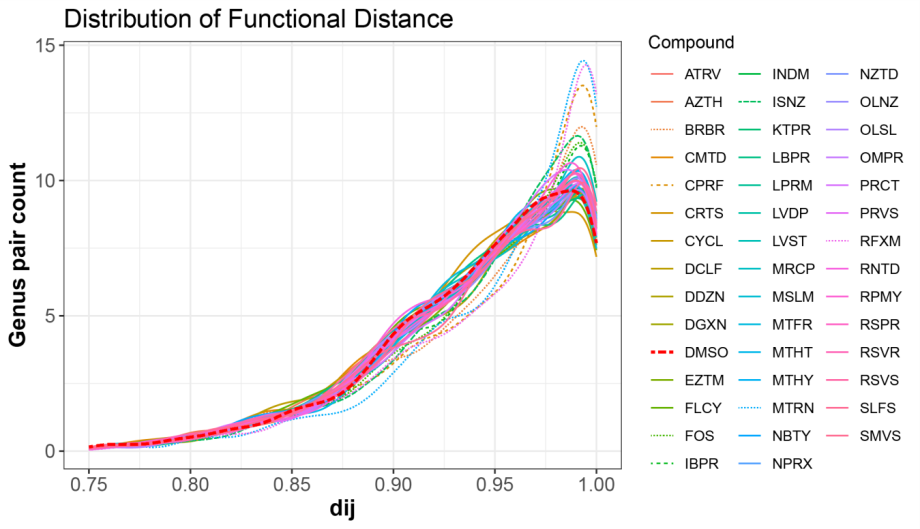

**Figure S9. Distribution of d*_ij_* values by compounds in the RapidAIM dataset.**

Each distribution line was plotted using the mean value across individual microbiomes (N=5) corresponding to the control (DMSO, red dashed line) or other compounds. Compounds shown in dashed lines, i.e. berberine (BRBR), ciprofloxacin (CPRF), fructo-oligosaccharide (FOS), ibuprofen (IBPR), isoniazid (ISNZ), metronidazole (MTRN) and rifaximin (RFXM) showed overall shifts in the distribution. Microbial genera comprising the top 95% protein biomass were involved in this figure.

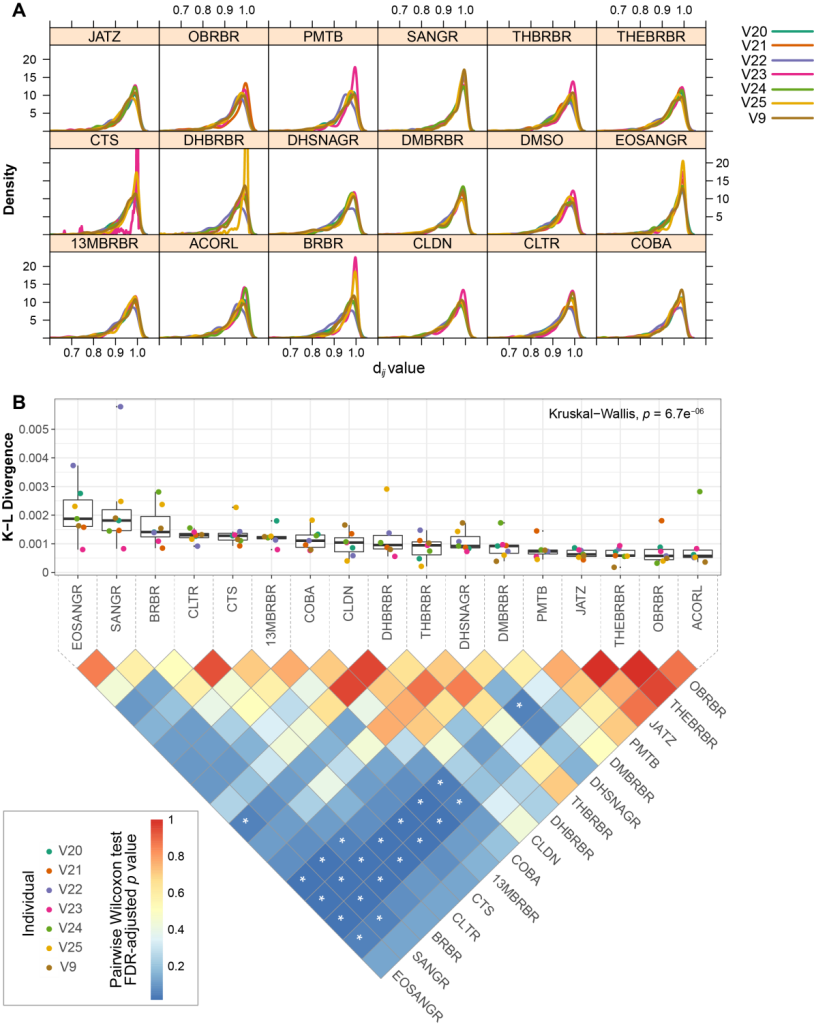

**Figure S10. Between-genera functional distances in the Berberine dataset.**

A. d*_ij_* distribution by different berberine analogues and by different individual microbiomes. B. K-L divergence between the d*_ij_* distribution in the control (DMSO) and that of the other compounds. Kruskal-Wallis test result indicated that overall the compounds had heterogeneous levels of K-L divergence with the DMSO. Between-compound comparisons of the K-L divergence values were performed by a Pairwise Wilcoxon Rank Sum Tests, “*” indicates statistical significance at the FDR-adjusted *p* < 0.05 level. The results were based on microbial genera of the top 95% overall protein biomass in the dataset.

Below are previews of Figures S11 and S12. They are provided as high-resolution PDF files separately.

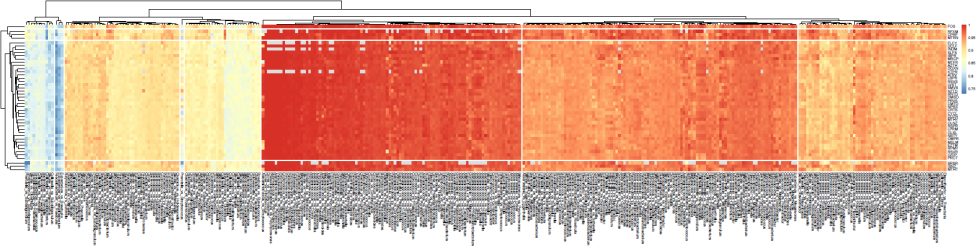

**Figure S11. Heatmap showing dij values between genera across compounds in the RapidAIM dataset**

Heat colors represent average d*_ij_* values in different individual microbiomes (N=5). Microbial genera comprising the top 95% protein biomass were used in this figure.

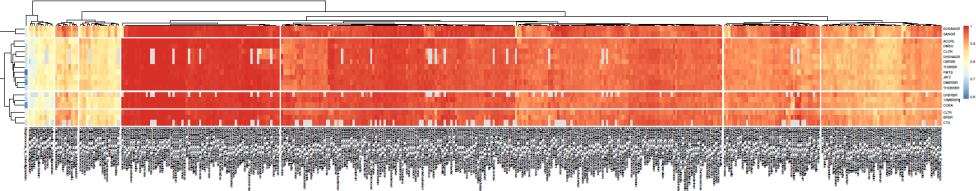

**Figure S12. Heatmap showing dij values between genera across compounds in the Berberine dataset**

Heat colors represent average d*_ij_* values in different individual microbiomes (N=7). Microbial genera comprising the top 95% protein biomass were used in this figure.

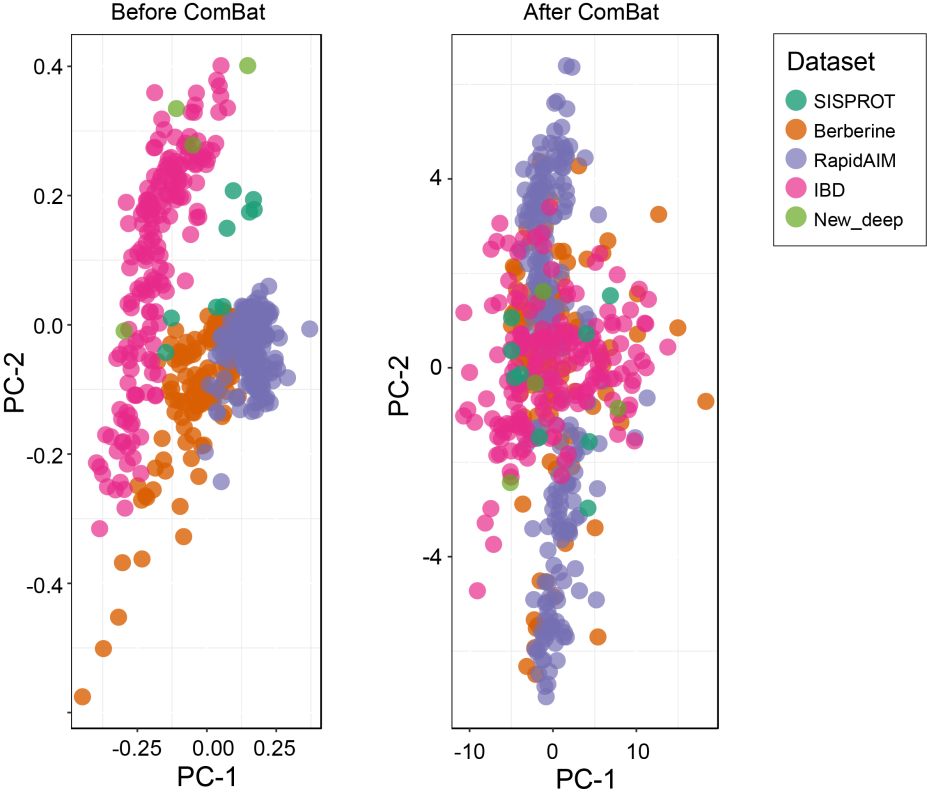

**Figure S13. Removing batch effects among datasets using ComBat**

Before the comparison of between-genera d*_ij_* values across metaproteomics datasets (**Figure 7**), an Empirical Bayesian approach (ComBat) was applied to remove batch effects between metaproteomics datasets with our Batch Effect Explorer <https://shiny1.imetalab.ca/batcheffect_explorer/>. PCA plots before and after the correction are shown here.
