## Supplementary notes for "Revealing Protein-Level Functional Redundancy in the Human Gut Microbiome using Ultra-deep Metaproteomics"

**Generating PCN from MaxQuant and MetaLab search results**

Through the Metapro-IQ workflow, we obtained the ProteinGroups.txt, and peptides.txt tables. The Protein groups table (generated by MaxQuant) contains information on the identified proteins, and identifiers of peptide sequence associated to each protein group. Through MetaLab, we further obtained MetaLab_peptide.xlsx table and function.csv tables. These tables are inter-connected through peptide sequences, peptide id numbers, and protein IDs. Therefore, we were able to match taxon and function by combining these result tables.

**Step-by-step workflow:**

A detailed step-by-step workflow is described below, as well as illustrated in Figures S14 and S15.

**Step 1.** The MetaLab_peptide.xlsx table (or set of tables) contain peptide sequences and the taxonomic matching according to the MetaLab pep2taxon database. And the peptide.txt file includes a column of unique peptide IDs for each peptide sequence. These two tables were first combined to generate a peptide_ID_to_taxon table.

**Step 2.** Each protein group in the ProteinGroup.txt table correspond to a series of peptide IDs, we are therefore able to link each protein group to the taxonomic information by querying these peptide IDs from the peptide_ID_to_taxon table. Protein group intensities were also kept in this table.

**Step 3**. The genus level information was summarized for each protein group to generate a ProteinGroup_genus_intensity table. Here, we approximately consider that the peptides corresponding to each protein group are derived from a same genus. We validated that this approach has a confidence of 98.4% at the genus level based on the ultra-deep metaproteomics dataset (**Supplementary Table S8**).

**Step 4.** Next, COG functions were taken from the top 1 protein in each protein group (function_top1 table). We validated that functions of proteins in each protein group have an agreement of 97.7% based on the ultra-deep metaproteomics dataset. In addition, top 1 protein in each protein group is considered the most confident protein identification, given by its number of identified peptides and E values.

**Step 5.** The function_top1 table was combined with the protein ProteinGroup_genus_intensity table to generate a ProteinGroup_function_taxon_intensity table.

**Step 6.** The ProteinGroup_function_taxon_intensity table can then be converted into PCN in the form of a bipartite network or an incidence matrix $\mathbf{P} = [P_{\mathrm{ia}}]$.

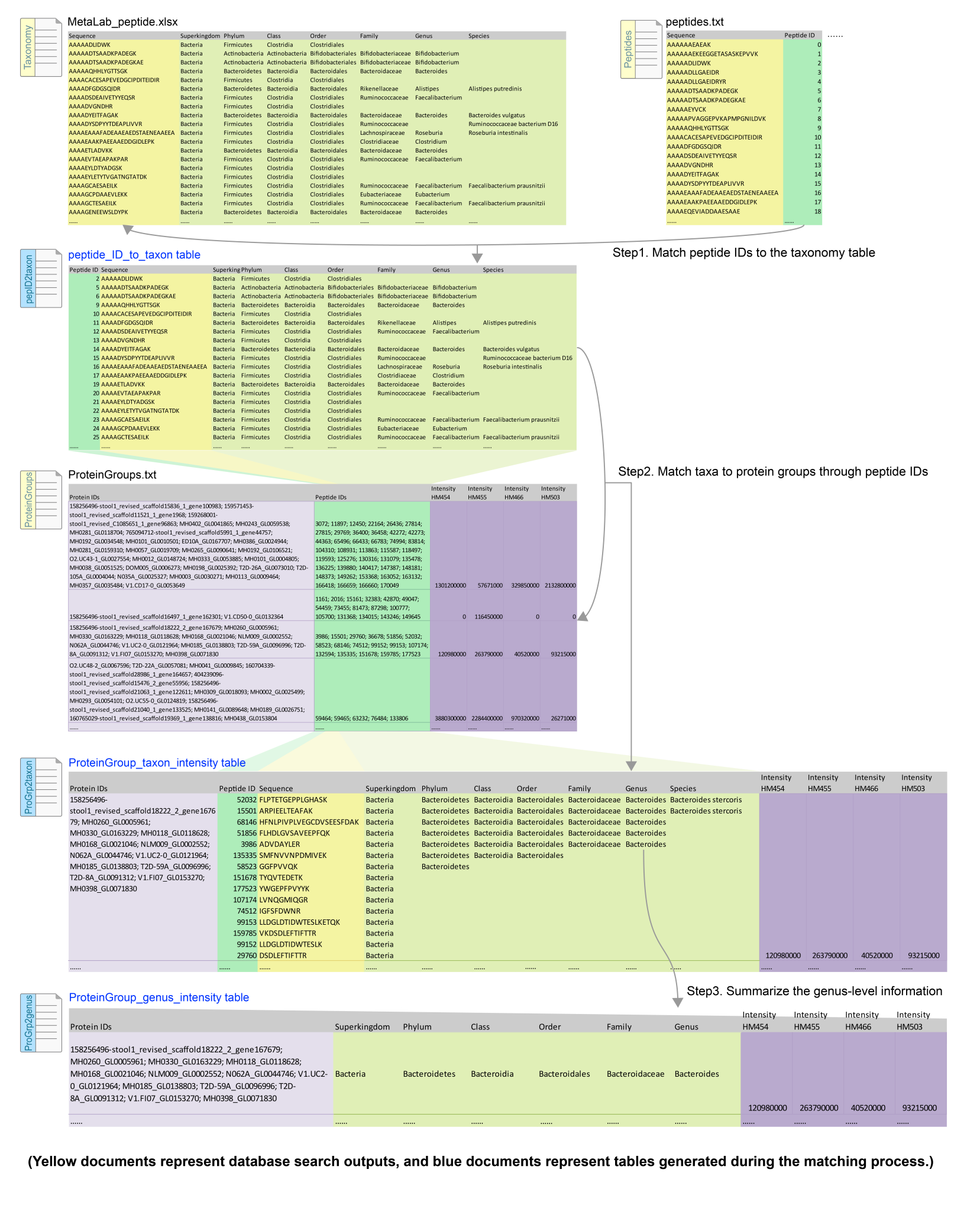

**Figure S14. Step-by-step workflow for PCN generation, part I.**

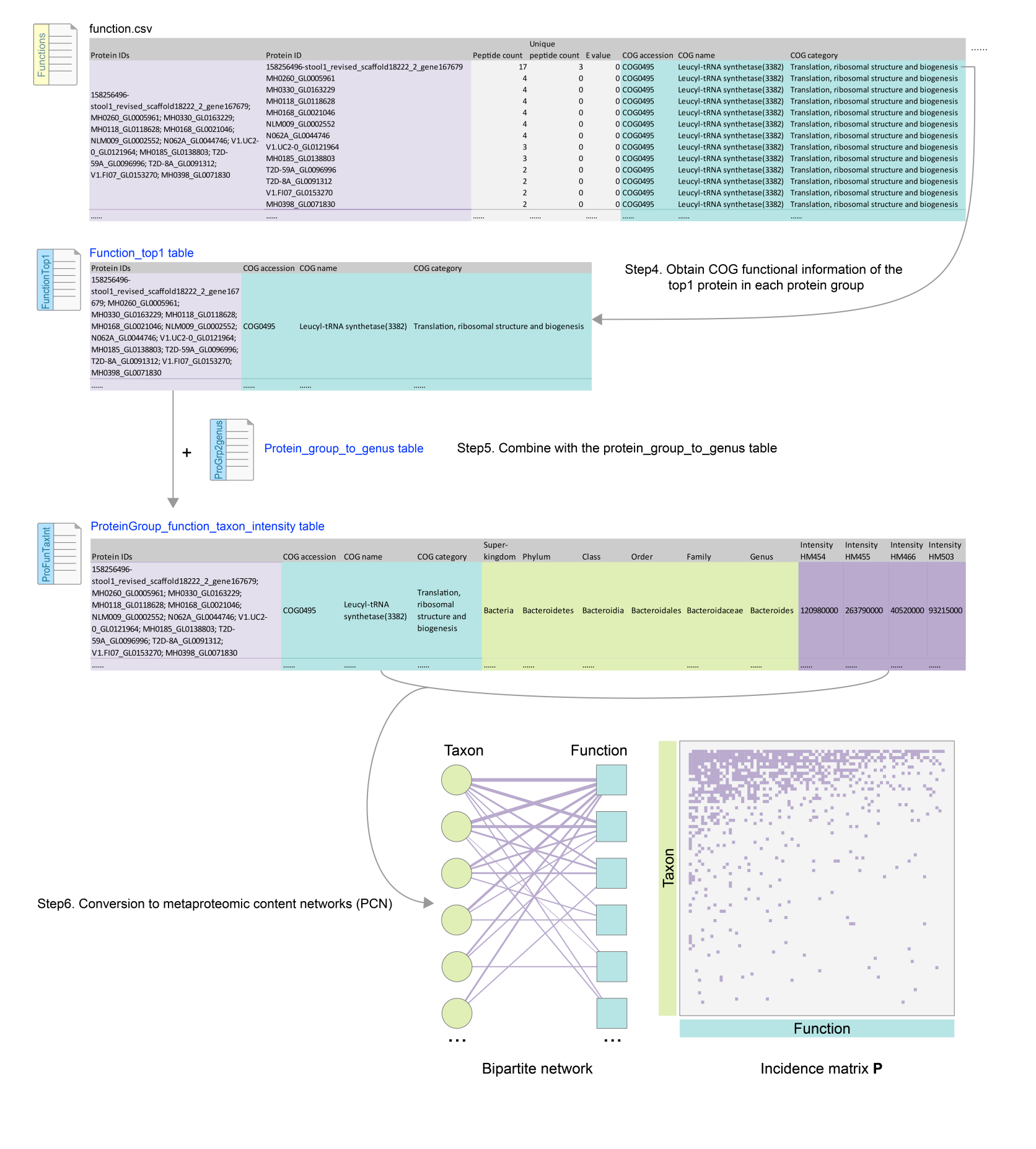

**Figure S15. Step-by-step workflow for PCN generation, part II.**

**Supplementary Table S8. Confidence of protein group-to-taxon matching**

| Taxonomic level: | Super-kingdom | | Phylum | Class | Order | Family | Genus | Species |
| --- | --- | --- | --- | --- | --- | --- | --- | --- |
| All unique pairs of matches at this level | | 46,592 | 44,894 | 41,269 | 41,016 | 29,425 | 26,748 | 15,477 |
| Protein groups matched to only one taxon at this level | | 46,553 | 44,491 | 40,900 | 40,778 | 28,855 | 26,322 | 15,104 |
| ProteinGroup% matched to a unique taxon at this level | | 99.9% | 99.1% | 99.1% | 99.4% | 98.1% | 98.4% | 97.6% |
| ProteinGroup% matched to more than one taxa at this level | | 0.1% | 0.9% | 0.9% | 0.6% | 1.9% | 1.6% | 2.4% |

Note: Numbers of matches were calculated using the ultra-deep metaproteomics dataset.
